# Post mortem mapping of connectional anatomy for the validation of diffusion MRI

**DOI:** 10.1101/2021.04.16.440223

**Authors:** Anastasia Yendiki, Manisha Aggarwal, Markus Axer, Amy F. D. Howard, Anne-Marie van Cappellen van Walsum, Suzanne N. Haber

## Abstract

Despite the impressive advances in diffusion MRI (dMRI) acquisition and analysis that have taken place during the Human Connectome era, dMRI tractography is still an imperfect source of information on the circuitry of the brain. In this review, we discuss methods for post mortem validation of dMRI tractography, fiber orientations, and other microstructural properties of axon bundles that are typically extracted from dMRI data. These methods include anatomic tracer studies, Klingler’s dissection, myelin stains, label-free optical imaging techniques, and others. We provide an overview of the basic principles of each technique, its limitations, and what it has taught us so far about the accuracy of different dMRI acquisition and analysis approaches.

## 1. Introduction

Diffusion MRI (dMRI) tractography was introduced two decades ago as a technique for reconstructing the trajectories of white-matter axon bundles by exploiting the anisotropy of water diffusion within these bundles (Mori et al., 1999; Jones et al., 1999; Conturo et al., 1999). Early applications of tractography demonstrated that it was generally able to reconstruct the large highways of the human brain, as they had been identified by prior anatomical studies (Catani et al., 2002; Wakana et al., 2004). However, this was only possible after extensive manual intervention, which was necessary to eliminate any paths that were reconstructed by tractography but that did not reflect true anatomy. The challenge in this process is that the true anatomy is not fully known – although we know the main pathways of the brain, we do not know the complete trajectories of all the small fiber bundles that comprise these pathways.

The fact that tractography is prone to errors is straightforward to establish using either simulated dMRI data (Côté et al., 2013; Daducci et al., 2014; Leemans et al., 2005; Neher et al., 2014; Maier-Hein et al., 2017), or real dMRI data collected from phantoms (Fieremans et al., 2008; Perrin et al., 2005; Poupon et al., 2008). In both of these scenarios, the ground-truth fiber geometry is known. Such studies are valuable for demonstrating the general limitations of tractography, and for comparing different tractography methods with respect to quantitative accuracy metrics. However, simulated and phantom data do not reflect the full complexity of brain circuitry. Thus the performance of a tractography method in such a setting cannot be used to determine which of the fiber bundles that this method reconstructs in a real brain are true and which are artifactual.

The past few years have seen a dramatic improvement in dMRI data quality, in large part due to technical advances spearheaded by the Human Connectome Project (Van Essen et al., 2013; Setsompop et al., 2013). These advances have increased the spatial and angular resolution, as well as the contrast-to-noise ratio, of the dMRI data that can be acquired routinely for *in vivo* studies. One of the main motivations behind this improvement is the expectation that higher quality of dMRI data, and specifically of data acquired with high *b*-values, should lead to more accurate tractography. The ultra-high *b*-values that can be achieved with these new technologies have been shown to reduce the uncertainty of probabilistic estimates of diffusion orientations (Setsompop et al., 2013) and to sharpen deterministic estimates of orientation distribution functions (Fan et al., 2014). These findings suggest improved ability to resolve crossing fiber bundles. However, not all fiber configurations in the brain can be modeled as crossings, and dMRI voxel sizes are still at a scale where multiple fiber configurations can lead to indistinguishable diffusion profiles. As a result, tractography is still imperfect. Therefore, the recent improvements in dMRI data quality have not obviated the need for validating the output of tractography algorithms; on the contrary, they have created the need for more sophisticated validation methods, capable of evaluating the fine-grained anatomy that can be captured by state-of-the-art dMRI.

Prior to the introduction of dMRI, all of our knowledge on the circuitry of the brain came from *post mortem* anatomical studies, using various techniques for dissection, tracing, histology, and microscopy. These are also the tools that we have at our disposal for *post mortem* validation of dMRI. The main focus of this review is connectional anatomy, hence we discuss techniques for validating the pathways that are output by tractography algorithms, or the local diffusion orientations that are the input to those algorithms. However, in many *in vivo* applications of tractography, the ultimate goal is to extract tract-specific biomarkers. In that sense, tractography is closely intertwined with dMRI microstructural modeling. Thus, we also survey the *post mortem* validation of microstructural parameters estimated from dMRI data.

For a precise, voxel-by-voxel comparison of white-matter circuitry and microstructure as obtained from dMRI and anatomy, both the dMRI and anatomical validation data should be collected from the same brain. Given the changes that a brain undergoes when it is excised and fixed, the dMRI scan should be collected after these procedures. Therefore, validation studies require expertise in both *ex vivo* dMRI scanning and anatomy. In section 2, we review the main methodological considerations for collecting *ex vivo* dMRI data. In section 3, we provide a brief overview of the evolution of anatomical studies, and specifically the techniques that can be used to obtain “gold standard” data for comparison to dMRI. We then discuss in more detail the validation of tractography (section 4), fiber orientations (section 5), and other microstructural parameters (section 6). We end with a discussion of open questions and future directions in section 7.

## 2. Ex vivo dMRI

*Ex vivo* dMRI acquisitions can harness the advantages of higher field and/or gradient strengths, more sensitive radiofrequency (RF) coils, and longer scan times. With tailored acquisition pulse sequences, isotropic spatial resolutions of a few hundred microns can be achieved for the whole brain and smaller specimens. This is almost an order of magnitude higher than typical *in vivo* resolutions of 1.5-2 mm. In recent years, *ex vivo* dMRI has emerged as a powerful tool to enable 3D mapping of human brain circuitry at mesoscopic scales, which is particularly important for comparison and validation of tractography methods using complementary *post mortem* modalities.

The main challenges for acquiring high-quality *ex vivo* dMRI data are the reduced *T*_2_ and reduced diffusivity of fixed tissue (D’Arceuil et al., 2007; Pfefferbaum et al., 2004; Roebroeck et al., 2019). While the former leads directly to loss of signal-to-noise ratio (SNR), the latter requires heavier diffusion weighting to offset the loss in contrast-to-noise ratio (CNR), but also contributes to further loss of SNR. Both of these factors need to be taken into account when designing optimal acquisition sequences and sampling schemes for *ex vivo* imaging. Below we discuss the main technical considerations for acquiring *ex vivo* dMRI data in the brain, and highlight the different acquisition sequences that can be used to achieve the high image quality and spatial/angular resolutions needed for tractography and microstructure validation studies.

Acquisition of *ex vivo* dMRI datasets often requires the use of tailored pulse sequences to combat the loss in SNR and CNR due to reduced *T*_2_ and diffusivity of fixed tissue. While 2D single-shot echo planar imaging (ss-EPI) sequences are most widely used for *in vivo* dMRI of the human brain, these are sub-optimal for *ex vivo* imaging, as the shorter *T*_2_ precludes the use of lengthy echo trains. Moreover, the higher *b*-values and stronger diffusion-weighting gradients used to offset the lower diffusivity can exacerbate eddy-current induced distortions when *k*-space is traversed in a single shot. Thus, multi-shot acquisition techniques and 3D echo trains are ideal for *ex vivo* dMRI. These techniques are not widely used *in vivo*, as they typically suffer from severe artifacts due to motion-induced phase errors across excitations. In *ex vivo* imaging, however, motion is not an issue.

Compared to 2D ss-EPI, 3D multi-shot or segmented EPI sequences allow for shorter effective echo times and thus higher SNR, and have been used for *ex vivo* dMRI of the whole human brain at 3 T (McNab et al., 2009; Miller et al., 2011; McNab et al., 2013) and the intact human brainstem at preclinical field strengths (Aggarwal et al., 2013). Multi-echo acquisitions involving repetitions of segmented EPI readouts at different echo times can be used to improve SNR (Eichner et al., 2020). On the other end of the spectrum, spin-echo (SE) readouts, which traverse a single line in *k*-space per excitation, have also been used for *ex vivo* dMRI (D’Arceuil et al., 2007; Dyrby et al., 2011; Guilfoyle et al., 2003; Modo et al., 2016). While these allow the highest anatomical fidelity with low geometric distortion, depending on the targeted spatial and angular resolution, they require prohibitively long scan times due to the relatively low SNR efficiency. At these lengthy scan times, problems with both magnetic field drift and specimen stability can become significant.

Diffusion weighted (DW) sequences with multiple RF-pulse echo trains, such as fast spin echo or gradient and spin echo, enable accelerated 3D imaging by factors of ∼4-12x as compared to DW-SE, while minimizing the distortion-related artifacts that DW-EPI sequences are prone to, and can therefore achieve combined high spatial and angular resolutions (Aggarwal et al., 2010; Tyszka and Frank, 2009). These may currently offer the best trade-off between SNR efficiency and image quality for smaller specimens, but require sophisticated schemes to correct for phase error induced artifacts, including but not limited to, the acquisition of navigator echoes or reference phase scans. Using volumetric excitations, the 3D *k*-space encoding can be further optimized to separate out eddy current and *T*_2_-decay effects on different *k*-space axes for techniques that combine gradient- and spin-echoes, thereby allowing dMRI data to be acquired with high SNR efficiency and reduced artifacts (Aggarwal et al., 2010). Such combined high spatial-angular resolutions are crucial, *e.g.,* for resolving fiber orientation distributions in the cortex (Aggarwal et al., 2015; Leuze et al., 2012).

In addition to optimized readout strategies to combat the effects of reduced *T*_2_, depending on the application at hand, modifications to the diffusion encoding may also be necessary. The *b*-values used for *ex vivo* dMRI need to be considerably higher than their *in vivo* counterparts, typically by factors of 2-4x, in order to achieve comparable diffusion contrast (Dyrby et al., 2011; Roebroeck et al., 2019; Schilling et al., 2017a). At such high *b*-values, eddy-current artifacts are particularly problematic, and can be further exacerbated when moving to higher field strengths. For micro-imaging applications (voxel sizes < 100 µm) at high field, bipolar diffusion-encoding gradients can be combined with multi-echo readouts to reduce the effects of eddy currents (Reese et al., 2003). As another alternative, DW steady-state free precession (SSFP) sequences, which can retain signal over multiple repetition intervals, have been shown to provide improved SNR efficiency with heavy diffusion weighting for whole brain *ex vivo* dMRI at 3 T and 7 T (Foxley et al., 2014; Miller et al., 2012). The slower *T*_1_ decay at higher field strengths can also be harnessed for strong diffusion weighting by employing stimulated-echo preparations, as shown for whole brain *ex vivo* dMRI at 9.4 T (Fritz et al., 2019).

*Ex vivo* acquisitions can also benefit from advances in gradient hardware or customized RF coils. Multi-channel coils, custom-built to closely fit the whole brain or smaller specimens, can allow increased reception sensitivity, thereby leading to a boost in the achievable SNR. For whole-brain imaging on human scanners, custom-built coils have been shown to lead to SNR gains of ∼1.6-2 fold as compared to standard *in vivo* head coils (Edlow et al., 2019; Roebroeck et al., 2015; Scholz et al., 2019). When combined with parallel transmit RF pulses for *B*_1_^+^ homogeneity, maximal SNR gain of as much as 5-fold was reported in the peripheral cortex using a custom-built cylindrical phased-array receive coil for the *ex vivo* occipital lobe at 9.4 T (Sengupta et al., 2018). Customized transmit/receive RF coils can be further combined with optimized acquisition pulse sequences such as DW-SSFP to achieve improved SNR for whole-brain *ex vivo* dMRI at submillimeter resolutions (Fritz et al., 2016). These advances in RF coils can enable the acquisition of much higher-quality dMRI validation data sets than what would be possible with standard, *in vivo* coils.

## 3. Anatomy: the gold standard for dMRI

The development of cellular and axonal markers at the turn of the 20^th^ century began the modern era of neuroanatomy. Two stains (Nissl and Golgi) provided the ability to visualize cell morphology, thus permitting the classification of cell types and the cytoarchitectonic organization of cortex. Degenerative stains made possible the visualization of myelinated axons, leading to a new understanding of connections between brain regions. These two anatomic subfields for understanding brain organization blossomed during the early 20^th^ century: cytoarchitectonics, which segmented the brain based on cortical layer cell morphology, and myeloarchitectonics, which classified cortical areas based on myelin distribution and fiber orientation through cortical layers (Vogt, 1903; Brodmann, 1909; Nieuwenhuys et al., 2015). Prior to the early 1950s, the only available anterograde tracer was the Marchi stain, which specifically marks degenerating myelin sheaths following well-placed lesions (Marchi and Algeri, 1885). However, as this method did not identify unmyelinated, thinly myelinated axons, or terminals, it was quite limited. While a reduced silver method did become available to visualize the axons themselves, shortly after, more sensitive tracing techniques were developed. These newer tracers relied on active neuronal transport, allowed the precise visualization of both axons and terminal fields, and became our most reliable source of information on connectional anatomy. We discuss these tracers as a validation tool for dMRI tractography in section 4.1. Although tracing experiments allow us to map axon bundles with high precision, they cannot be conducted on human subjects. The only alternative in the human brain is to perform blunt dissection on fixed *ex vivo* specimens. The methodology that is used for this purpose to this day was devised by Joseph Klingler in the 1930s. It involves loosening the structure of fixed brain tissue, and particularly fiber bundles, by freezing and thawing the tissue (Klingler, 1935; Ludwig and Klingler, 1956; Klingler and Goor, 1960). We discuss Klingler’s dissection method as a technique for validating dMRI tractography in section 4.2.

In tracing and blunt dissection studies, the goal is to follow a group of axons from its origin to its terminations. When the goal is to visualize all the axon bundles that intersect in a given brain location, for comparison with a fiber orientation distribution (FOD) computed from dMRI data at a certain voxel, we need other techniques. In neuroanatomy, this type of information is usually obtained by processing tissue sections with histological stains for myelin. The precursor to such stains was developed in 1873 by Camillo Golgi, one of the pioneers of modern-day histology, who perfected a silver-staining method that he coined “the black reaction” (Glickstein 2006). The stain had affinity to only a few neurons, but stained their structure - soma, dendrites and axon - in its entirety. We discuss histological stains as a method for validating fiber orientations in section 5.1.

In the past decade, novel optical imaging techniques have been adopted to visualize fiber orientations in the brain. These include label-free methods, like polarization microscopy and optical coherence tomography, which rely on intrinsic tissue contrast instead of staining. Another novel approach is clearing, *i.e.,* rendering tissue transparent, after which the tissue is treated with fluorescent dyes and imaged with fluorescence microscopy. These methods are far less developed than traditional histological techniques, but they hold great promise as the digital neuroanatomy tools of the future. We discuss these methods for validating fiber orientations in section 5.2.

Studies that have performed comparisons of dMRI tractography and anatomic tracing are summarized in Table 1. Studies that have compared dMRI orientation estimates and microscopic measurements of fiber orientations in the same sample are summarized in Table 2.

**Table 1.**
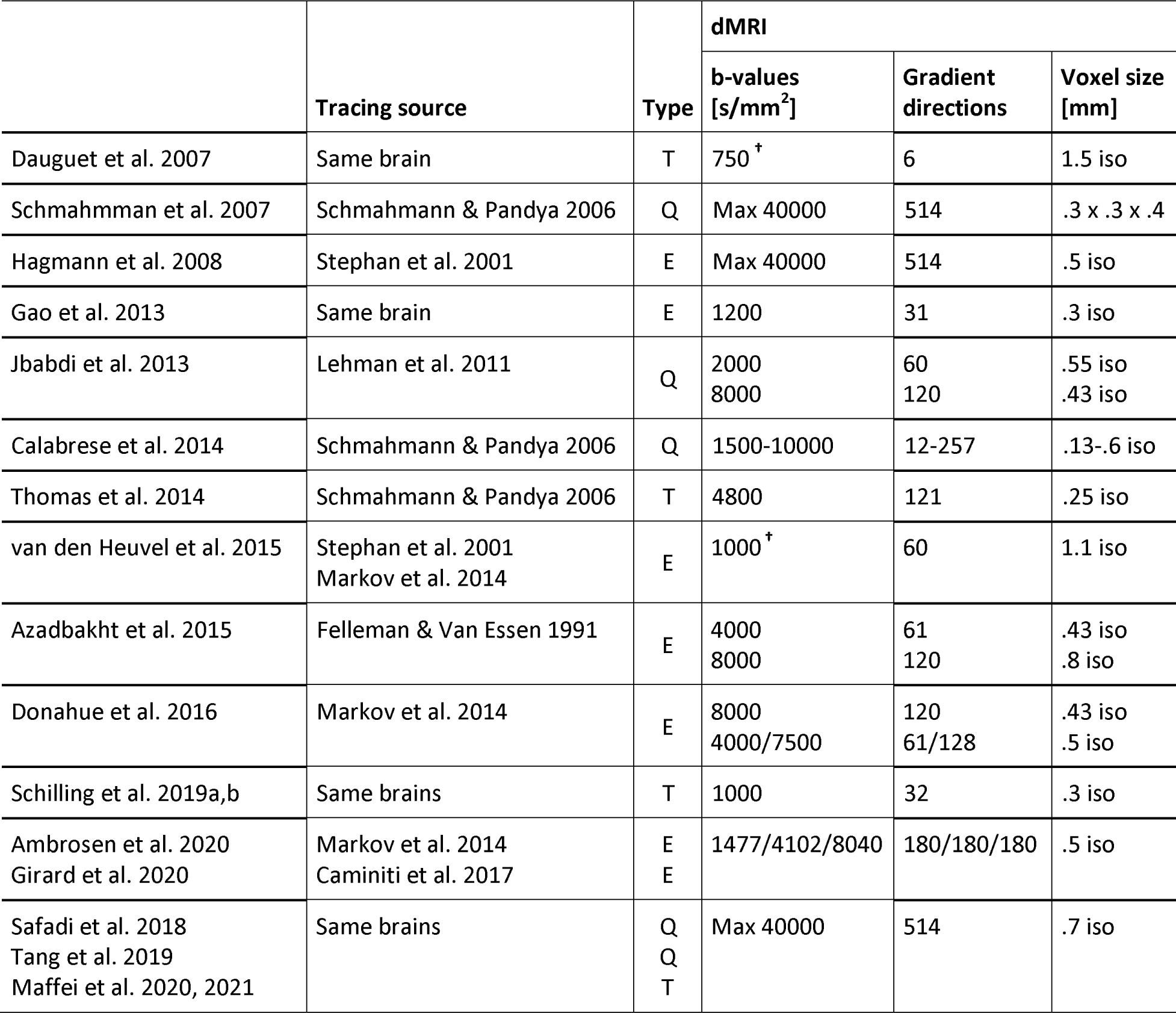
Summary of literature: validation of tractography. Studies that have performed comparisons of dMRI tractography and anatomic tracing in non-human primates. We list the source of the tracer data (from a previously published database or from the same brain as the dMRI scan), type of comparison (E: endpoints; Q: qualitative; T: trajectory), and dMRI acquisition parameters for each study (“iso” indicates isotropic voxels).^†^: These b-values were collected *in vivo;* hence they are not directly comparable to the b-values of the other studies.

|  | Tracing source | Type | dMRI |  |  |
| --- | --- | --- | --- | --- | --- |
|  |  |  | b-values [s/mm<sup>2</sup>] | Gradient directions | Voxel size [mm] |
| Dauguet et al. 2007 | Same brain | T | 750<sup>†</sup> | 6 | 1.5 iso |
| Schmahmman et al. 2007 | Schmahmann & Pandya 2006 | Q | Max 40000 | 514 | .3 x .3 x .4 |
| Hagmann et al. 2008 | Stephan et al. 2001 | E | Max 40000 | 514 | .5 iso |
| Gao et al. 2013 | Same brain | E | 1200 | 31 | .3 iso |
| Jbabdi et al. 2013 | Lehman et al. 2011 | Q | 2000<br>8000 | 60<br>120 | .55 iso<br>.43 iso |
| Calabrese et al. 2014 | Schmahmann & Pandya 2006 | Q | 1500-10000 | 12-257 | .13-.6 iso |
| Thomas et al. 2014 | Schmahmann & Pandya 2006 | T | 4800 | 121 | .25 iso |
| van den Heuvel et al. 2015 | Stephan et al. 2001<br>Markov et al. 2014 | E | 1000<sup>†</sup> | 60 | 1.1 iso |
| Azadbakht et al. 2015 | Felleman & Van Essen 1991 | E | 4000<br>8000 | 61<br>120 | .43 iso<br>.8 iso |
| Donahue et al. 2016 | Markov et al. 2014 | E | 8000<br>4000/7500 | 120<br>61/128 | .43 iso<br>.5 iso |
| Schilling et al. 2019a,b | Same brains | T | 1000 | 32 | .3 iso |
| Ambrosen et al. 2020<br>Girard et al. 2020 | Markov et al. 2014<br>Caminiti et al. 2017 | E<br>E | 1477/4102/8040 | 180/180/180 | .5 iso |
| Safadi et al. 2018<br>Tang et al. 2019<br>Maffei et al. 2020, 2021 | Same brains | Q<br>Q<br>T | Max 40000 | 514 | .7 iso |

**Table 2.** Summary of literature: validation of fiber orientations. Studies that have compared dMRI orientation estimates in post-mortem brain tissue to independent measurements of fiber orientations from a reference modality in the same sample. We list the reference modality, type of tissue, and dMRI acquisition parameters for each study (“iso” indicates isotropic voxels).

|  | Reference modality |  | Tissue type | dMRI |  |  |
| --- | --- | --- | --- | --- | --- | --- |
|  |  |  |  | b-values [s/mm<sup>2</sup>] | Gradient directions | Voxel size [mm] |
| Leergaard et al. 2010 | Myelin stain | 2D | Rat | Max 30452 | 514 | .265 iso |
| Choe et al. 2012 |  | 2D | Owl monkey | 1309 | 21 | .3 iso |
| Seehaus et al. 2015 |  | 2D | Human | 3000 | 60 | .34 iso |
| Schilling et al. 2017 |  | 2D | Macaque | 6000 | 101 | .3-.8 iso |
| Schilling et al. 2016 | DiI | 3D | Squirrel monkey | 3200/6400 | 30/90 | .4 iso |
| Schilling et al. 2018 |  | 3D | Squirrel monkey | 3000/6000<br>9000/12000 | 100/100<br>100/100 | .2 x .2 x .4 |
| Mollink et al. 2017 | PLI | 2D | Human | 5000 | 120 | .4 iso |
| Hannsen et al. 2019 |  | 2D | Human | 4000 | 256 | .5 iso |
| Wang et al. 2014 | PSOCT | 2D | Human | 4032 | 20 | .3 iso |
| Jones et al. 2020 |  | 2D | Human | Max 40000 | 514 | .25 iso |
| Stolp et al. 2018 | CLARITY | 3D | Mouse | 1500 | 42 | .125 x .125 x .2 |
| Leuze et al. 2021 |  | 3D | Human | 2000 | 447 | 1.0 iso |

## 4. Validation of tractography

### 4.1 Anatomic tracing

#### 4.1.1 Methodology

Anatomic tracing studies in non-human primates (NHPs) allow a direct visualization of the wiring of the brain, including cells of origin, axon trajectories, and terminals. This level of detail allows the appreciation of the full complexity of brain connections, and makes tracer studies an anatomic “gold standard” for interpreting and validating the indirect measurements of the wiring of the brain that we obtain with dMRI tractography.

The first tracers that relied on active neuronal transport were preferentially transported anterogradely (e.g., tritiated amino acids) or retrogradely (e.g., Horse radish peroxidase-HPR conjugated to wheat germ agglutin-WGA). Soon other molecules followed with intrinsic fluorescence and with better sensitivity, which could be further increased with immunohistochemical processing. Tracer molecules are microinjected in the brain region of interest, taken up into the cell body, and transported to the terminal fields (anterograde tracers) or taken up by the terminal fields and transported to the cell body (retrograde tracers). Different tissue processing protocols are applied to visualize the labeled axons, terminal fields and cell bodies, depending on the specific molecule injected. Nonetheless, they all provide the ability to directly visualize and quantify the labeled cells, axons, and their pathways through the white matter and terminal fields.

Details on the surgery, perfusion, and histological processing involved in tracer studies are provided in Haber et al (2006) and Lehman et al (2011). Briefly, animals receive an injection of one or more of the following antero-grade/bidirectional tracers: Lucifer yellow, fluororuby, or fluorescein conjugated to dextran amine (LY, FR, or FS; 40–50 nl, 10% in 0.1M phosphate buffer (PB), pH 7.4; Invitrogen); *Phaseolus vulgaris*-leucoagglutinin (PHA-L; 50 nl, 2.5%; Vector Laboratories); or tritiated amino acids (100 nl, 1:1 solution of [3H] leucine and [3H]-proline in dH2O, 200 mCi/ml, NEN). Twelve to fourteen days after surgery, animals are again deeply anesthetized and perfused with saline followed by a 4% paraformaldehyde/1.5% sucrose solution in 0.1 M PB, pH 7.4. Brains are postfixed overnight and cryoprotected in increasing gradients of sucrose (10%, 20%, and 30%).

Serial sections of 50 μm are cut on a freezing microtome into 0.1 M PB or cryoprotectant solution. One in eight sections are processed free-floating for immunocytochemistry to visualize the tracers. Tissue is incubated in primary anti-LY (1:3000 dilution; Invitrogen), anti-FS (1:1000; Invitrogen), anti-FR (1:1000; Invitrogen), or anti-PHA-L (1:500; EY Laboratories) in 10% NGS and 0.3% Triton X-100 (Sigma-Aldrich) in PB for 4 nights at 4°C. Following extensive rinsing, the tissue is incubated in biotinylated secondary antibody followed by incubation with the avidin-biotin complex solution (Vectastain ABC kit, Vector Laboratories). Immunoreactivity is visualized using standard DAB procedures. Sections are mounted onto gel-coated slides, dehydrated, defatted in xylene, and coverslipped with Permount. Sections for autoradiography are mounted on chrome alum gelatin-coated slides and defatted in xylene overnight. Slides are dipped in Kodak NTB2 photographic emulsion and exposed for 4 – 6 months at 4°C in a light-tight box and then developed in Kodak D19 for 2.5 min., fixed, washed, and counterstained with cresyl violet. Each slide that has been processed to visualize a given tracer is annotated under the microscope to outline the fiber bundles as they travel from the injection site. The slides are co-registered to each other and the outlines assembled across slices to produce a 3D reconstruction of the bundles.

#### 4.1.2 Comparison to dMRI

There are three main approaches to comparing anatomic tracing to dMRI tractography, and they differ in terms of the information that is extracted from the tracer data. These approaches are: *(i)* comparing only the areas of origin and termination of the bundles, *(ii)* comparing the topography of the bundles as they travel through the white matter, and *(iii)* comparing the exact position of the bundles throughout their trajectory.

##### (i) Comparison of areas of origin/termination

This is the approach taken by most prior studies that have compared anatomic tracing to dMRI tractography. Here the information on brain connections is reduced to a “connectivity matrix” that indicates which brain region is connected to which. These general connection patterns are highly reproducible between individual brains, hence it is reasonable to perform such a comparison using dMRI data and tracer experiments from different brains. This approach has been popular because it can take advantage of existing data on the “connectivity matrix” of the brain from prior NHP tracer studies, such as those included in the CoCoMac or other published databases (Felleman and Van Essen, 1991; Stephan et al., 2001; Markov et al., 2014). The accuracy of tractography is evaluated based on the frequency of the following outcomes: dMRI tractography detects a connection between a pair of brain regions that is also shown to be connected based on tracer experiments (true positive); it misses such a connection (false negative); or it detects a connection between a pair of regions that is not shown to be connected based on the tracer data (false positive).

Several studies have generated area-to-area connectivity matrices using tractography in dMRI scans of NHP brains and compared them to existing collections of NHP tracer data (Hagmann et al., 2008; van den Heuvel et al., 2015; Azadbakht et al., 2015; Donahue et al., 2016; Ambrosen et al., 2020). Despite the variety in dMRI acquisition and analysis methods, as well as the tracer databases used by these studies (see Table 1), there is considerable agreement in receiver operating characteristic (ROC) analyses. For the most part, for a specificity around 60%, tractography achieves sensitivity in the 60%-70% range in most studies (Hagmann et al., 2008; Azadbakht et al., 2015; Donahue et al., 2016; Ambrosen et al., 2020). A study that performed this analysis with a variety of more modern tractography algorithms, found that the sensitivity at the same level of specificity edged somewhat higher, in the 60%-80% range (Girard et al., 2020). One of the previous studies examined the effect of dMRI acquisition parameters on the accuracy of the connectivity matrix (Ambrosen et al., 2020). A notable finding was that increasing the *b*-value from 1477 to 8040 led to indistinguishable ROC performance.

Analyses of correlation between the connectivity matrices obtained from dMRI tractography and tracer databases show less agreement than ROC analyses. Reported correlation coefficients range from r=.25 (van den Heuvel et al., 2015) to r=0.59 (Donahue et al., 2016). Beyond the NHP literature, one study relied on a database of tracer injections in ferrets (Delettre et al., 2019). After regressing out the distance between cortical areas, correlation between connectivity matrices obtained from dMRI tractography in a ferret scan and from the tracer database was much lower for deterministic tractography (r=.36-.40; not statistically significant) that probabilistic tractography (r=.54-.77; statistically significant). The only study that performed this type of correlation analysis using dMRI and tracer data from the same animal, a squirrel monkey, showed much higher correlation of area-to-area connectivity matrices, which ranged from r=.73 to r=.93, depending on the dMRI analytic approach (Gao et al. 2013). This may suggest that using dMRI and tracer data from different animals underestimates the accuracy of tractography. However, differences in analysis steps between these studies make it difficult to draw a definitive conclusion on this point.

##### (ii) Comparison of topographies

This approach requires tracer data that go beyond those widely available in databases, as it examines the trajectories of the axon bundles through the white matter and not just their areas of origin/termination. Tracer studies are an excellent source of information on the topographic organization of axon bundles, *i.e.,* the relative positions of smaller bundles as they travel through a large white-matter pathway such as the internal capsule, corpus callosum, or cingulum bundle (Lehman et al., 2011; Heilbronner et al., 2014). Fig. 1 shows an example of axon bundles projecting from three cortical areas into the internal capsule. The dorsal-to-ventral topography of these cortical areas is preserved by the corresponding bundles as they travel through the internal capsule. This is revealed by NHP tracing experiments and replicated with tractography in both *ex vivo* NHP dMRI and *in vivo* human dMRI.

**Figure 1.**
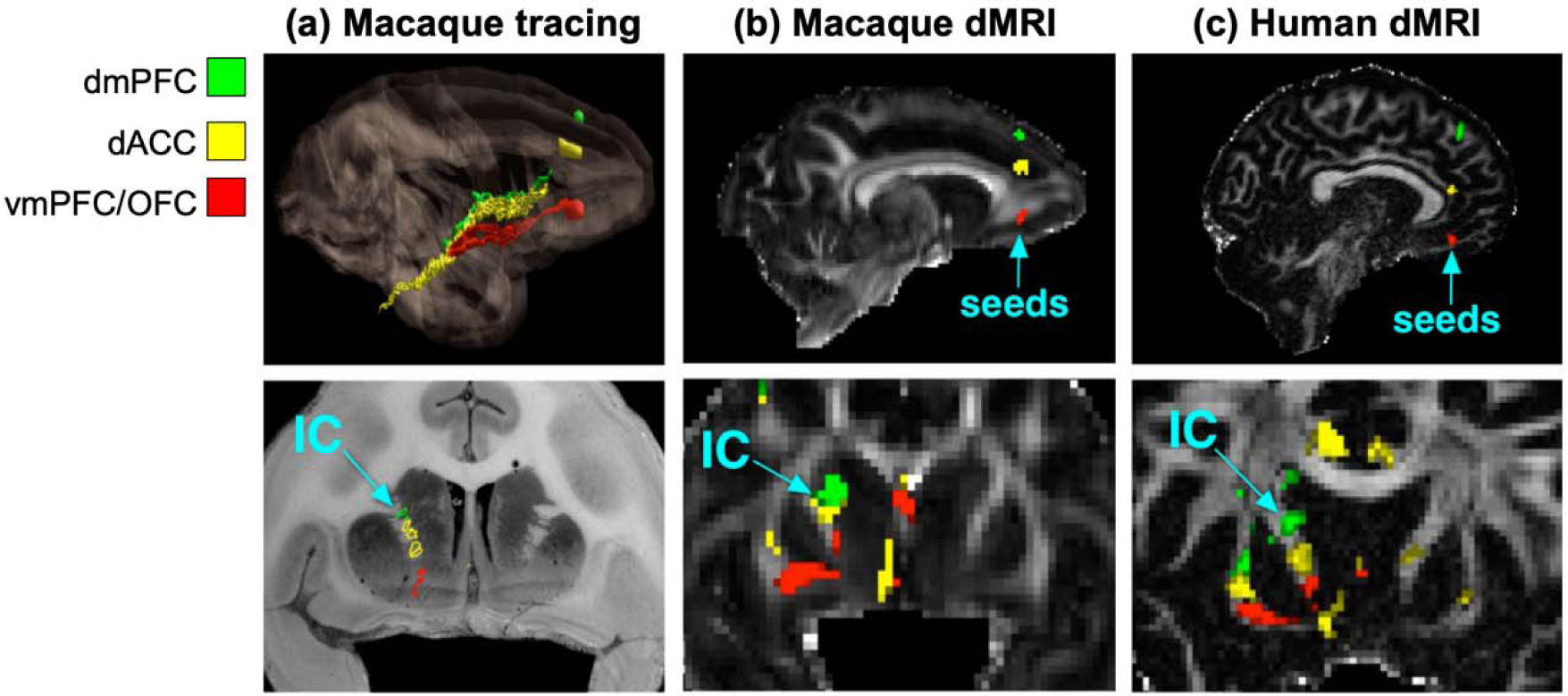
Topographies of axon bundles shown with anatomic tracing vs. dMRI. The projections of the dorsomedial prefrontal cortex (dmPFC), dorsal anterior cingulate cortex (dACC), and ventromedial prefrontal cortex (vmPFC) / orbitofrontal cortex (OFC) follow a dorsal-to-ventral topographic organization in the internal capsule (IC). This is shown by placing tracer injections in each of the three areas in NHP **(a)** and replicated by seeding dMRI probabilistic tractography in the three areas, in both *ex vivo* NHP data **(b)** and *in vivo* human data **(c)**. (Adapted from Safadi et al., 2018.)

While dMRI tractography is known since its inception to reconstruct the large highways of the brain, replicating the fine topographies of smaller axon bundles within them is a far more challenging task, and hence a more rigorous testbed for the accuracy of tractography methods. Comparisons of topographies in NHP tracing, NHP ex vivo dMRI, and human in vivo dMRI show that many of them can be replicated by tractography in both NHPs and humans (Safadi et al., *2018).* Importantly, this work demonstrates that, while the absolute coordinates of small axon bundles in template space are inconsistent across individual subjects, the relative positions of these bundles with respect to each other are highly consistent. This finding has two key implications. First, it suggests that validation of dMRI tractography in terms of such organizational rules could reasonably be performed using dMRI and tracer data from different brains. Second, it demonstrates that it is more reliable to define white-matter bundles in terms of their relative positions with respect to each other and to their surrounding anatomy than it is to define them in terms of absolute coordinates in a template space.

##### (iii) Voxel-by-voxel comparison

This is a much more detailed level of validation, where the precise route of the axon bundles through the brain is compared between tractography and tracing on a voxel-wise basis. While comparisons of dMRI and tracing at the two previous levels (areas of origin/termination or topographic organization) are valuable for assessing *how often* tractography errors occur, voxel-wise comparisons are the only way to determine exactly *where* the errors occur. This is critical for revealing which fiber configurations are the most common failure modes of tractography algorithms, and therefore identifying ways to improve these algorithms. For this type of validation study, tracer and dMRI data must come from the same brain. (See Grisot et al., 2021 for an analysis of intra- vs. inter-individual variability.) Even if the general patterns of connectional anatomy are similar across brains, it is unlikely that image registration will lead to perfect voxel-wise alignment of all bundles, and especially of the small groups of axons that are labeled by a tracer injection.

Some of the early studies that compared tractography to anatomic tracing in terms of the trajectory of the bundles through the white matter did not have dMRI and tracer data from the same animals (Schmahmman et al., 2007; Jbabdi et al., 2013; Calabrese et al., 2014). Hence, the comparisons were performed in a qualitative manner, ensuring that tractography could replicate the route of bundles that had been previously described in the anatomy literature. Another study superimposed a set of illustrations of tracer maps from a previously published collection to a dMRI scan from a different animal (Thomas et al., 2014). The authors evaluated deterministic and probabilistic tractography, but only performed a full ROC analysis on the latter. For data points where the deterministic and probabilistic approaches have approximately the same true or false positives, the latter appears to outperform the former.

An early study that performed anatomic tracing and dMRI scanning in the same NHP brain was able to achieve a Dice coefficient (i.e., overlap) between tracing and tractography as high as 0.75, despite being limited by low spatial and angular resolution (Dauguet et al., 2007). Another dataset that included anatomic tracing and dMRI in the same animals was used to compare a wide range of tractography algorithms, including as part of the VoTEM challenge (Schilling et al., MRI 2019a,b). These data confirm a result from (Thomas et al., 2014), that the default thresholds for probabilistic tractography operate in a regime with higher true and false positives than the default thresholds for deterministic tractography.

A more recent study used dMRI and tracer data from the same animal, collected with a more advanced dMRI acquisition protocol that the previous studies (Grisot et al., 2021). This allowed the comparison of multiple q-space sampling schemes (Cartesian, single-, and multi-shell). A full ROC analysis revealed that probabilistic tractography consistently achieved higher true positive rates than deterministic tractography, when the two were compared at the same false positive rates, for a variety of orientation reconstruction methods and sampling schemes. The maximum *b*-value had only a modest impact on the accuracy of tractography. Finally, the voxel-wise comparison of tractography and tracer data in the same brain revealed which axonal configurations led to errors consistently, across dMRI acquisition and analysis methods. These common failure modes of tractography involved geometries such as branching or turning, which are not modeled well by conventional crossing-fiber reconstruction methods. The same data were used as the training case for the IronTract challenge, with an additional injection in a different brain serving as the validation case (Maffei et al., 2020, 2021). The first round showed that, when tractography pipelines were optimized to achieve maximum accuracy for one injection site, they were generally much less accurate for the other site. Two teams were the exception to this rule, exhibiting robustness across the two seed regions (Maffei et al., 2020). In the second round, all teams used the pre- and post-processing methods that had been used by these two top teams. This increased both accuracy and robustness between seed areas for most other teams. One of the two post-processing strategies involved a simple Gaussian filter and the other applied a set of *a priori* anatomical inclusion masks. Remarkably, Gaussian filtering led to similar improvements in the accuracy of tractography methods as the use of a priori anatomical constraints (Maffei et al., 2021).

#### 4.1.3 Limitations

The first, and perhaps most important, limitation of tracer studies is that they can only be performed in animals. The benefit of performing these studies in NHPs is that holomologies between the monkey and human brain have been studied extensively. Despite the expansion of the frontal lobe in humans, similarities in position, cytoarchitectonics, connections, and behavior indicate that the circuitry is relatively comparable (Uylings and van Eden, 1990; Petrides, 1994; Petrides and Pandya, 1994; Ongür and Price, 2000; Chiavaras and Petrides, 2000; Petrides and Pandya, 2002; Petrides et al., 2012). Thus, a possible route for using the knowledge generated by NHP tracer studies to validate dMRI tractography in humans is via the first two approaches described in the previous section, *i.e.,* validation in terms of end regions or topographies. Of course, it is possible that a difference between the tracing in NHPs and dMRI tractography in humans may be due to a true inter-species difference. This is where dMRI in NHPs can serve as a crucial stepping stone. When the NHP tracing disagrees with dMRI in both NHP and human, this is more likely to be due to an error in dMRI than an inter-species difference (Safadi et al., 2018). Furthermore, if the bundles reconstructed with dMRI tractography in humans become more similar to those seen in the NHP tracing when dMRI data quality improves, we can be more confident about using NHP tracer data and across-species homologies to evaluate tractography in human dMRI (see Fig. 2 for an example).

**Figure 2.**
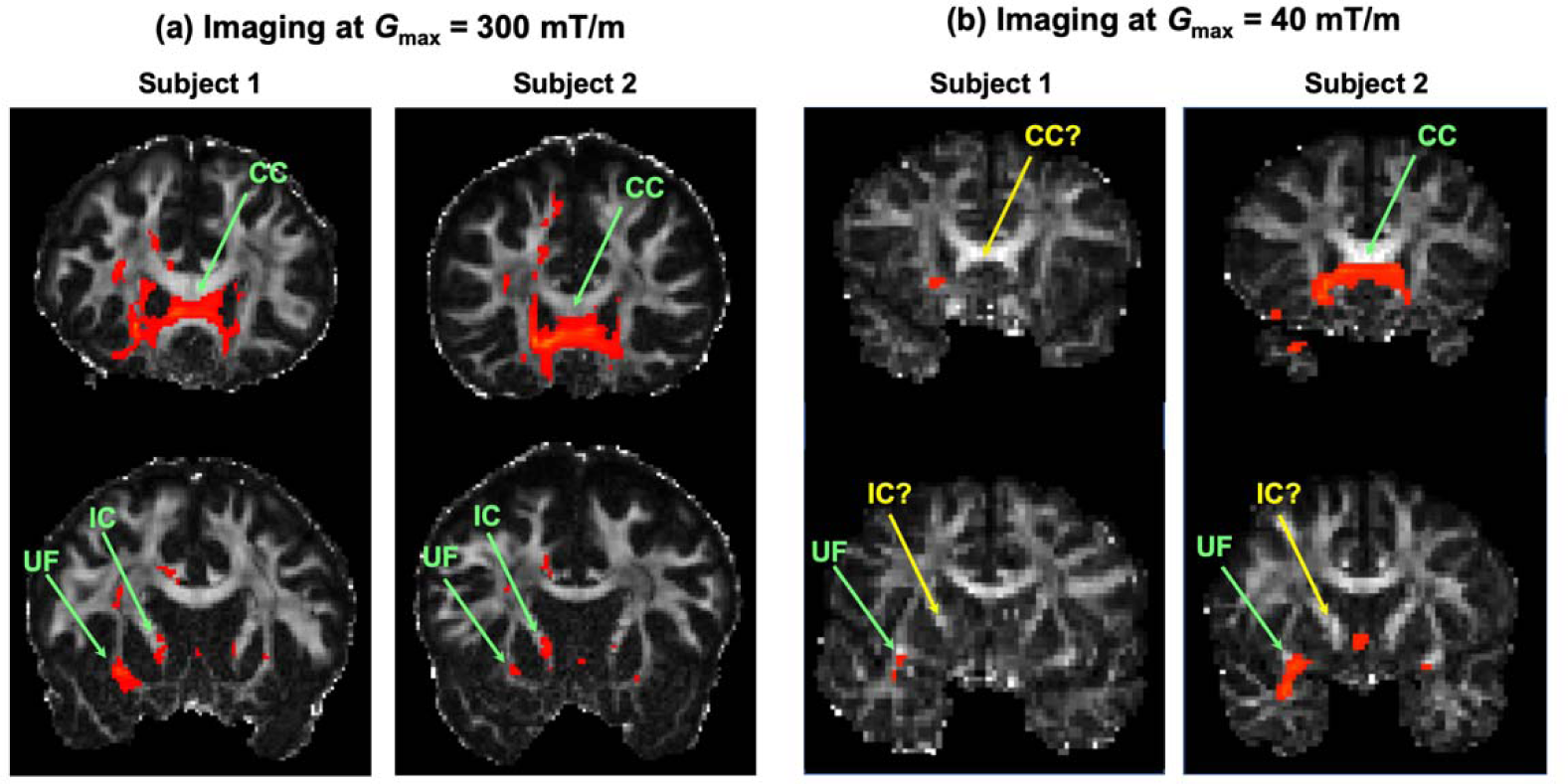
Improved *in vivo* human dMRI shows greater agreement with NHP anatomic studies. Probabilistic tractography from a seed region in the frontal pole is shown for two volunteers imaged on the *Connectom* scanner (a), and two imaged on a scanner with conventional gradients (b). Based on NHP anatomic studies (Lehman et al., 2006), we expect to find frontal pole projections in the corpus callosum (CC), internal capsule (IC), and uncinate fasciculus (UF), among other pathways. These are present consistently in the *Connectom* data, while many are missing in the conventional data. True positives are marked with green arrows; false negatives with yellow arrows. When discrepancies between human dMRI and NHP tracing disappear as dMRI data quality improves, these discrepancies are more likely to be dMRI errors than true inter-species differences.

Tracers suffer from a variety of other limitations. Conventional tracers can be taken up by fibers of passage and the exact area of axonal uptake at the injection site can be difficult to determine. In addition, inconsistency in uptake and transport can result in variable quality between injections. These conventional tracers have now largely been replaced by viral tracers, optogenetic methods, etc., for rodent work. Unfortunately, some of these more modern anatomic methods that circumvent these issues are difficult and costly to carry out in old-world NHPs.

Finally, as is the case for all methods that rely on histological processing (see section 5.1 for a detailed discussion), distortions due to sectioning and staining can interfere with the alignment of histological sections. However, tracer studies have the advantage that the same axons are labeled by the tracer across slices, and this provides a visual aid in ascertaining that slices have been aligned correctly. In the future, the acquisition of more data with tracing and dMRI in the same brains, and the development of algorithms for automated annotation and analysis of axon bundles from tracer data will be critical for harnessing the full potential of anatomic tracing as a tool for the validation of dMRI tractography.

### 4.2 Gross dissection

#### 4.2.1 Methodology

The tracer studies described in the previous section are not applicable to human subjects. Klingler’s dissection technique is the only means available to anatomists for following fiber bundles in *ex* vivo human brains. Detailed descriptions of this technique are provided in the reviews by Wysiadecki et al. (2019) and Dziedzic et al. (2021). In short, the brain is fixed in a 10% formalin solution. It must remain in fixative for at least 2 months, and suspension prevents it from deforming during the fixation process. The meninges are carefully removed. The brain is frozen at −10 to −15°C for one week and then thawed by washing with running water for one day. The brain is immersed again in new formalin and the freezing-thawing procedure can be performed a second time. Klingler and colleagues recommended freezing the specimens before dissection, because they thought that the formalin solution did not penetrate the myelinated nerve fibers fully, resulting in higher formalin concentrations between the fibers. When the specimens are frozen, formalin ice crystals form between the nerve fibers, expanding and separating them. The freezing process facilitates the dissection of fine fiber bundles in particular (Türe et al. 2000). Soft wooden spatulas are used to peel away the fiber bundles along the anatomic planes. When fibers are too thin for the eye to see, an operating microscope can be used, with a ×6 to ×40 magnification.

Protocols vary among different publications on Klingler’s technique. Differences are found in the perfusion method (whole body perfusion or brain perfusion via the vertebral and carotid arteries), freezing time, frequency of the freezing procedure, temperature and formalin percentage. Türe et al. (2000) and de Castro et al. (2005) provide clear descriptions of Klingler’s method. Best practices from these two reports are combined with our own experience in the detailed protocol of Arnts et al (2013).

Türe et al. (2000) and de Castro et al. (2005) describe the removal of the pia mater, arachnoid membrane, and vessels of the specimens using the operative microscope. However, with some experience it is possible to remove the membranes and vessels without a microscope (Arnts et al. 2013). Both Türe et al. (2000) and de Castro et al. (2005) also recommend that the brains be stored in a refrigerator at −10 to −15 degrees Celsius and placed in fresh 10% formalin solution for 8–10 days. Freezing time differs in Pujari et al. (2008), who report a freezing time of 14 days. Castro et al. (2005) is the only one to repeat the freezing periods and freeze the brains in a formalin solution. For the dissection step itself, all authors use wooden spatulas to tease out the white matter tracts by blunt dissection. Lawes et al. (2008) use spatulas of 3 mm or 6 mm width. Most authors also use an operation microscope. As with brain preparation, the dissection technique differs between publications. Some report on a cortex-sparing dissection technique, where not all cortex is scraped away (Martino et al., 2011). Regardless of the specifics of the technique, however, knowledge of white matter anatomy is crucial for Klingler’s dissection. For a review, see Schmahmann (2008) and Mandonnet (2018).

With the introduction of dMRI and tractography, white matter dissection has regained interest in the neuroimaging and neuroanatomical community. Klingler’s dissection is still practiced to date, not only for research on white-matter anatomy, but also for developing surgical approaches (Türe et al. 2000; Baydin 2016), and for the preparation of educational specimens (Arnts et al. 2013).

#### 4.2.2 Comparison to dMRI

Ideally, when white-matter tracts dissected with Klingler’s method are compared with those reconstructed by dMRI tractography, both dissection and dMRI scanning should be performed in the same brain. Ex vivo dMRI scanning followed by Klingler’s dissection was performed on two vervet monkey brains to investigate the existence of the inferior fronto-occipital fasciculus, and found evidence for it from both techniques (Sarubbo et al., 2019). Dissections of the optic radiations in ex vivo human brain specimens have been compared to dMRI tractography of the same tract in a different set of ex vivo human brains (Nooij et al., 2015).

In most cases, however, the results of Klingler’s dissection have been compared to tractography results from *in vivo* dMRI scans of a different set of subjects (Lawes et al., 2008; Martino et al., 2011; Martino et al., 2013; Goryainov et al., 2017; Latini et al., 2017; Maffei et al., 2018; Briggs et al., 2019,2020,2021; Flores-Justa et al., 2019; Bernard et al., 2020; Li et al., 2020; Shinohara et al., 2020; Egemen et al., 2021; Weiller et al., 2021). In those cases, only qualitative comparisons between tractography and dissection have been possible. Sometimes, differences in reconstructions of the same tract across different subjects is reported not only with tractography, but also with dissection. For example, Goryainov et al. (2017) performed dissections of the superior longitudinal fasciculus in 12 brain specimens and found it to be subdivided into two components in 10 of the specimens and three components in the remaining two specimens. In some cases, Klingler’s dissection on its own is able to demonstrate shortcomings of tractography. For example, a study that performed dissections on fetal brains revealed that several of the main white-matter pathways were present, at earlier gestational stages than what had been previously shown by dMRI studies (Horgos et al., 2020). This suggests that the inability of tractography to reconstruct these pathways *in utero* is likely to be due to technical limitations of fetal MRI scanning.

Zemmoura et al. (2014) used a laser scanner to capture the surface and texture of a post mortem brain repeatedly as it underwent Klingler’s dissection. They then used this reconstruction to align ex vivo MRI and dissection results from the same brain. They reported a total accuracy for the method in the order of 1 mm. This included the accuracy of the surface-based, cross-modal registration, the deformation of the specimen during dissection, and the distance between two consecutive surface acquisitions with the laser scanner. This technique was a step toward a quantitative comparison of dMRI tractography with dissection.

#### 4.2.3 Limitations

Although the precision of Klingler’s dissection method is sometimes questioned, an electron microscopy study of fiber microstructure at different stages during the freezing, thawing, and dissection showed that, while the procedure destroyed most extra-axonal structures, it preserved myelin sheaths (Zemmoura et al., 2016).

Nonetheless, Klingler’s technique has several shortcomings as a tool for the validation of dMRI tractography. First, it cannot chart the complete trajectory of fibers, as it cannot follow them into the grey matter. Thus, the exact termination of axon bundles in the cortex or subcortical structures cannot be determined. Furthermore, the technique has difficulty following fibers through areas of dense crossing, as in the centrum semiovale, where commissural, projection, and association fibers intersect. Klingler’s technique is suitable for dissecting large pathways, but not for distinguishing small axon bundles that travel through them or for mapping the topographic organization of smaller axon bundles within the large pathways.

There are several steps in the preparation of the brain that are key for a successful comparison between dissection and dMRI: early extraction of the brain after death, a long fixation time at a low concentration of formalin, and a long period of freezing (Zemmoura et al., 2014). Any deviation from best practices can lead to errors. If the brain is to also undergo MRI scanning, a short post mortem interval (within 24 hours of death) is key for obtaining good MR contrast. This limits the availability of suitable specimens. Finally, dissection with Klingler’s technique is time consuming and requires extensive neuroanatomical expertise. As a result, most studies are performed on a small number of specimens. Given the substantial inter-individual variability in the geometry of the human brain, and the fact that the dissection is typically compared to dMRI tractography from a different set of brains, it is difficult to ascertain if any discrepancies between dissection and tractography results are due to errors of tractography or true individual differences.

## 5. Validation of fiber orientations

### 5.1 Histological stains

#### 5.1.1 Methodology

Histology is used extensively in research to describe tissue microanatomy and in clinical practice for the diagnosis and staging disease through tissue biopsies. Excised tissue is sectioned into thin slices and chemically stained. Histology slides can be digitized using high-throughput slide scanners, producing 2D images with micron or sub-micron resolution. In this section, we focus on the histological stains that are most relevant to visualizing fiber orientations: myelin and silver stains. While these techniques have been mainly used to validate fiber configurations locally within a histological section, it is also possible to apply 3D reconstruction to these sections and use them to follow fiber bundles through the brain (Mollink et al, 2016; Alho et al., 2021).

##### Myelin stains

The modified **Heidenhain-Woelcke stain** (Bürgel et al., 1997; Holl et al., 2011) was designed to provide contrast to distinguish between densely packed axon bundles, such as the optic radiation in post mortem human brain. Traditional myelin stains do not provide contrast to distinguish between different axon bundles. The modification of this protocol inactivates the chromatogen complexes in the thinnest myelin sheaths. This produces a graded reduction in myelin staining in white matter that appears to be proportional to the amount of myelination (Bürgel et al., 1997, 1999, 2006). This myelin stain is based on the presence of lipoproteins in the neuroceratine skeleton of myelin heaths (Bürgel et al., 1997).

**Luxol Fast Blue (LFB)** stains myelin blue. Carriel et al. (2017) described three histochemical methods for the staining of myelin, suitable for formalin-fixed and paraffin-embedded tissue. One method is the conventional luxol fast blue (LFB) method. The Klüver-Barrera stain (Klüver and Barrera, 1953) combines LFB with cresyl violet, which stains cell nuclei pink or purple. The Luxol Fast Blue MBS salt stains the myelin sheet blue. In ethanol solution, the salts’ anion—copper phthalocyanine—binds to the cationic elements of the tissue. A differentiation step with lithium carbonate can remove weakly bound anions from gray matter, while the stronger ionic bonds with the lipoproteins in the myelin sheet remain intact. The conventional LFB method is especially useful in the central nervous system, but its weak contrast limits its use in peripheral nerves (Carriel et al., 2017). As can be seen in **Fig. 3**, LFB stains the whole subcortical white matter blue and is not able to distinguish the various white-matter fasciculi or U-fibers. However, the stria of Gennari in the primary visual cortex (V1), where myelinated fibres are abundant, can be identified with LFB (Kleinnijenhuis, 2014).

**Figure 3.**
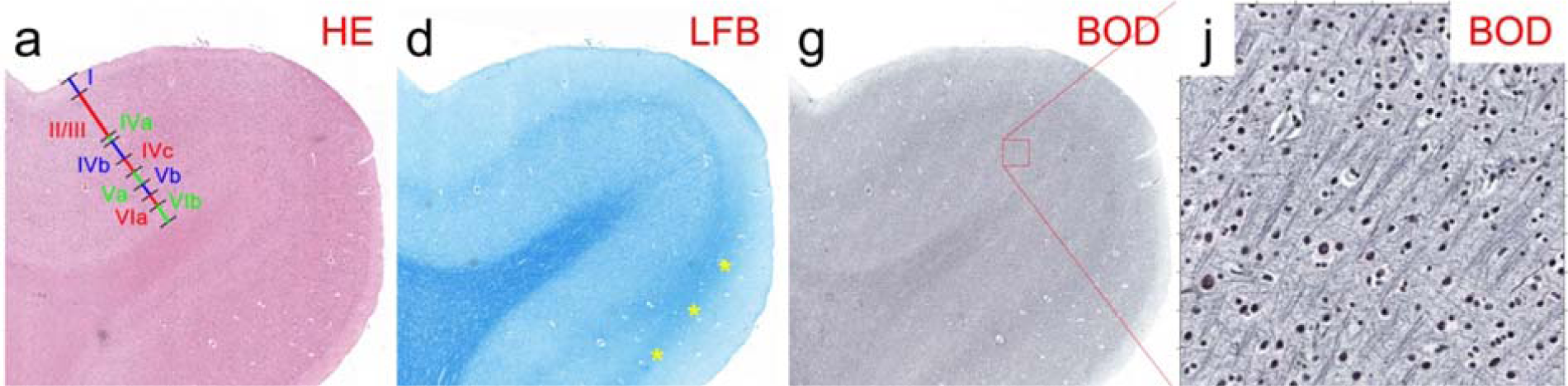
Histological sections from paraffin-embedded human brain samples around the calcarine sulcus containing the stria of Gennari, stained with: **a:** Hematoxylin & Eosin (HE; 5 μm). Cortical layers are identified as I-VIb. **d:** Luxol Fast Blue (LFB; 5 μm). **g:** Bodian (BOD; 6 μm). **j:** Detail of g. (Adapted fro Kleinnijenhuis, 2014.)

##### Silver stains

**Bodian** is a silver stain that stains the neurofilaments in all compartments of the neuron, i.e., axons, dendrites, and cell body. It stains all neurites black (Bodian, 1936). It is a argyrophilic silver staining method, i.e., ions from a silver proteinate solution bind to neurofilaments in the cytoskeleton (Gambetti et al., 1981) and, with the help of a reducing reagent (hydroquinone), metallic silver particles are formed. The silver is then replaced by metallic gold from a gold chloride solution (using oxalic acid as reductant), a process known as toning. In the Bodian method, the silver proteinate is combined with metallic copper. The silver protein oxidizes the metallic copper, and both are reduced onto the section by hydroquinone. During the toning stage, both silver and copper are replaced by gold, resulting in more intense staining (Uchihara 2007).

A comparison of LFB and Bodian stains is shown in Fig. Y. The LFB-stained section has a particularly high fiber volume fraction in the white matter and stria of Gennari, where myelinated fibres are abundant. In the infragranular layers, LFB staining is much denser when compared to the supragranular layers. The Bodian stain shows modest contrast between gray and white matter. However, more detail can be seen intracortically with Bodian than with LFB. In the supragranular layers, which appear rich in unmyelinated fibers, Bodian staining is relatively denser than LFB.

Another silver staining method for myelin is the **Gallyas stain** (Gallyas, 1979). It is known to label myelin well and produce clearly interpretable histology images but also to be less consistent than other protocols, i.e., it will often show a substantial amount of variance within and between tissue probes (Seehaus et al., 2015).

##### Other techniques for labeling myelin

The **black-gold stain** (Schmued and Slikker, 1999) and **Heidenhain-Woelcke stain** (Romeis, 1989) have been previously used to map myeloarchitecture in the human cortex (Eickhoff et al. 2005). **Immunohistochemistry** has also been used for the identification of myelin, using antibodies against **myelin basic-protein** (MBP; Kuhlmann et al., 2017) or **proteolipid-protein** (PLP; Mollink et al., 2017). For PLP staining in Mollink et al (2017), the tissue was paraffin embedded and sections were counterstained with haematoxylin (HE).

##### Lipophilic dyes

Lipophilic carbocyanine dyes, such as DiI and DiO, spread by lateral diffusion within the cell membrane (Honig and Hume 1989a,b). These dyes can be used as post mortem tracers in the human brain, where the use of active-transport tracers (see section 4.1.1) is not an option. However, tracing with DiI in human post mortem material is challenging. Although the speed of diffusion increases with temperature, the propagation of these dyes in the tissue is very slow. Hence, they can only be used to trace short-range connections, e.g., to follow fibers within a histological section. Tangential spread of about 8 mm was shown with DiI in the hippocampal formation (Tardif and Clarke, 2001). The technique is also appropriate for showing dendritic alterations in cortical neurons (Thal et al., 2008). A recent study described a modification of the method, where ethanol-dissolved DiI was used to achieve much faster diffusion than conventional DiI in fixed human brain (Sivukhina et al., 2021).

#### 5.1.2 Comparison to dMRI

In the following we discuss studies that used histological stains to validate fiber orientation estimates. These studies varied widely in terms of their dMRI acquisition protocols. They ranged from around 20 diffusion-encoding directions with a low b-value, in which case only single-tensor fitting could be performed, to 514 directions with a Cartesian-grid sampling scheme, in which case full orientation distribution functions (ODFs) could be fit to the data.

Most of these studies relied on myelin stains. In-plane (2D) fiber orientations were extracted from the histological sections by manual labeling (Leergaard et al., 2010), Fourier analysis (Choe et al., 2012), or structure tensor analysis (Seehaus et al., 2015; Schilling et al., 2017b). Reported angular errors include 5.7° for ODFs in a two-way fiber crossing area of rat brain (Leergaard et al., 2010) and less than 10° for tensors in a single-fiber area of owl monkey brain (Choe et al., 2012). The angular error of tensors in human M1 was found to increase from 10° to 40° as the fractional anisotropy (FA) decreased from 0.3, indicating more coherent fibers, to 0.05, indicating less coherent fibers due to more fanning or crossing (Seehaus et al., 2015).

A potential complication when evaluating dMRI orientation reconstruction methods in terms of their angular error is that the methods that tend to detect more fiber populations, i.e., produce ODFs with more peaks (some of which may be spurious), also tend to have lower angular errors. Thus it is important to also validate the number of peaks. A study that compared the number of fiber populations extracted from myelin-stained histological sections to that detected by dMRI found that this number increased as the voxel size decreased (Schilling et al., 2017b). This may seem counter-intuitive at first, if one expects that increasing the spatial resolution should lead to more voxels with a single fiber population. However, the multiple, distinct fiber populations found in smaller voxels can merge into single but more dispersed fiber populations as the voxel size increases. This is an example where the ground-truth fiber configurations obtained from post mortem microscopy can challenge the simplistic assumptions that are made in the development of dMRI analysis techniques.

An alternative to myelin staining is to stain sections of white matter with DiI or DiA (fluorescent lipophilic dyes) and subsequently image them with confocal microscopy. In this case, axonal orientations can be obtained with structure tensor analysis (Budde and Frank 2012). A benefit of this approach is that it can be extended to compute not only in-plane but 3D orientations from each histological section (Khan et al. 2015). A study that applied this technique to the squirrel monkey brain reported angular errors of 11.2° for tensors and 6.4° for fiber ODFs in areas with a single fiber population, and 10.4°/11.6° for fiber ODFs in the primary/secondary orientation of crossing-fiber areas (Schilling et al., 2016). Fiber ODFs has less than 20% success rate at resolving fiber populations that crossed at angles smaller than 60°. A follow-up study, which examined a greater variety of single-shell dMRI protocols and reconstruction methods for resolving crossing fibers, reported a median angular error of around 10° in voxels with a single fiber population and 11°/16° in the primary/secondary peak of voxels with multiple fiber populations (Schilling et al., 2018). There was little change in the angular error when the b-value increased from 6,000 to 12,000 or when the number of diffusion-encoding directions increased from 64 to 96.

#### 5.1.3 Limitations

Histological processing involves a laborious sequence of steps, which include embedding, sectioning, mounting on glass slides, staining, and cover-slipping. These procedures require considerable expertise and can be error-prone. Notably, the tissue undergoes non-linear physical deformations (warping and tearing) during sectioning. Such deformations make it difficult to align consecutive histological sections to each other and to the target dMRI volumes. Complex registration frameworks have been proposed to reduce such distortions and improve the alignment of histological and MRI data (Lebenberg et al., 2010; Choe et al., 2011; Adler et al., 2014; Majka and Wójcik 2016; Iglesias et al., 2018). A common approach is to align each distorted histological section to an undistorted blockface photograph taken before the section is cut, and then align the stacked sections from the blockface space to the MRI volume. This involves a series of 2D (slice-to-slice) registrations, followed by a 3D (volume-to-volume) registration. The registration algorithms used in each step involve several free parameters (deformation model, image similarity metric, regularization metrics, multi-resolution scheme, interpolation method, etc.) Each of these has its own trade-offs and can thus introducing subjectivity. At every step of the process, neuroanatomical expertise is key for evaluating the quality of the registration.

In addition to deformations due to sectioning, the dehydration that the tissue undergoes during staining can lead to tissue shrinkage (Wehrl et al., 2015; Williams et al., 1997). This may introduce discrepancies in fiber orientations pre- vs. post-staining, which may compound the estimated angular errors of dMRI orientations. It is important, however, to remember the difference in scale between the two modalities. Errors at the microscopic scale of the histological data may not have a significant impact on computations performed at the mm scale of the dMRI data.

Another possible concern arises when only in-plane fiber orientations are available from the histological data. In this case, the 3D diffusion orientations are projected onto the plane of the histological section and the angular error is computed in 2D. One may question whether such a comparison is as informative as one that uses the full 3D fiber orientations. Encouragingly, 2D and 3D angular errors reported in the literature (see previous section) are comparable. Finally, validation studies that rely on histology require that fiber orientations be computed, e.g., by Fourier or structure tensor analysis, from the stained and scanned sections. These image processing steps can be an additional source of errors. The following section describes optical imaging techniques that can measure fiber orientations directly, as an intrinsic contrast of the tissue.

### 5.2 Optical imaging

#### 5.2.1 Methodology

The 21^st^ century has seen a renaissance in light microscopy applications in neuroscience, driven by a combination of advances in tissue preparation and labeling methods, automation for faster image acquisition, and increased computational power. Here we focus on the latest optical imaging techniques that are specifically targeted at the visualization of axonal orientations.

*Polarization microscopy* and *polarization-sensitive optical coherence tomography* belong to the family of *label-free methods.* That is, they do not use exogenous contrast agents, such as stains, dyes, or tracers. Instead, they rely exclusively on a contrast mechanism that is intrinsic to the tissue. Specifically, these methods rely on *birefringence*, a property of anisotropic tissues. The use of polarization microscopy as a tool for visualizing myelinated fibers in both normal and pathological nervous tissue has been described in numerous studies for more than a century (e.g., Ehrenberg, 1849; Brodmann, 1903; Fraher et al., 1970; Miklossy et al., 1987). The basic principle is to generate polarized light, pass it through a thin brain section, and measure alterations in the polarization state of the light. This generates contrast between structures with different optic axis orientations. In white matter, the optic axis of the tissue is defined by the orientation of myelinated axons. For polarization microscopy, most microscopes operate in transmission mode, with few exceptions working in reflection mode (Takata et al., 2018).

The most prominent method for polarization microscopy is *3D Polarized Light Imaging (3D-PLI),* introduced by Axer M et al. (2011a,b). It uses a physical model to estimate the orientation of the optic axis of the underlying tissue directly from the measured sinusoidal signal. Fiber orientations can be not only reconstructed within sections, but also followed across sections. The orientation vectors can then be displayed as color-coded fiber orientation maps (FOMs) or combined over a neighborhood of microscopic-resolution voxels to compute a fiber orientation distribution (FOD) (Axer et al., 2016; Alimi et al., 2020). When the neighborhood over which the FOD is computed represents a mesoscopic-resolution dMRI voxel, this FOD can serve as the ground truth for validating the FOD or ODF obtained from dMRI.

Brain preparation is crucial in polarization microscopy, as the organization of the lipid bilayers composing the myelin sheaths has to be preserved. The tissue is immersed in a buffered solution of formaldehyde (4% in Axer M et al., 2011a,b; 7.7% in Henssen et al., 2019) for several days to months depending on the sample size, and in a cryoprotectant such as glycerin or sucrose. Sectioning is done with a cryostat microtome. In principle, PLI could be performed on sections of any thickness. In practice, however, due to constraints related to cryo-sectioning and handling large-area sections, the typical thickness is 50-100 μm, *i.e.,* 2-5 times thicker than the sections used for conventional histology (*e.g.,* Amunts et al., 2020). Polarizing microscopes can achieve in-plane resolutions ranging from 100μm (Axer H et al., 2011), 64 μm (Axer M et al., 2011a), 4 μm (Mollink et al., 2017), to 1.3 μm (Reckfort et al., 2015), and fields of view ranging from a few mm^2^ to a whole human brain section (up to 300 cm^2^). Large areas of interest are imaged tile-wise and the tiles are assembled by stitching.

The use of PLI to image brain tissue has been demonstrated in human (Axer et al., 2011a,b; Caspers et al., 2015; Mollink et al., 2017; Zeineh et al., 2017; Henssen et al., 2019), seal (Dohmen et al., 2015), rat (Schubert et al., 2016, 2018), pigeon (Herold et al., 2018; Stacho et al., 2020), and vervet monkey (Takemura et al., 2020). Examples of the detailed visualizations of cortical and white-matter fiber architecture that can be achieved by PLI are shown in Fig. 4. Research is ongoing on several extensions to this technology. Transmitted light intensity measurements have recently been shown to differentiate between areas with low fiber density and in-plane crossing fibers vs. areas with out-of-plane fibers, thus removing a potential confound for PLI (Menzel et al., 2020). Finally, *scattered light imaging (SLI)* is a new, label-free technique that can resolve crossing fibers within a microscopic-scale (6.3) voxel and can be integrated into a polarization microscope (Menzel et al., 2020, 2021).

**Figure 4.**
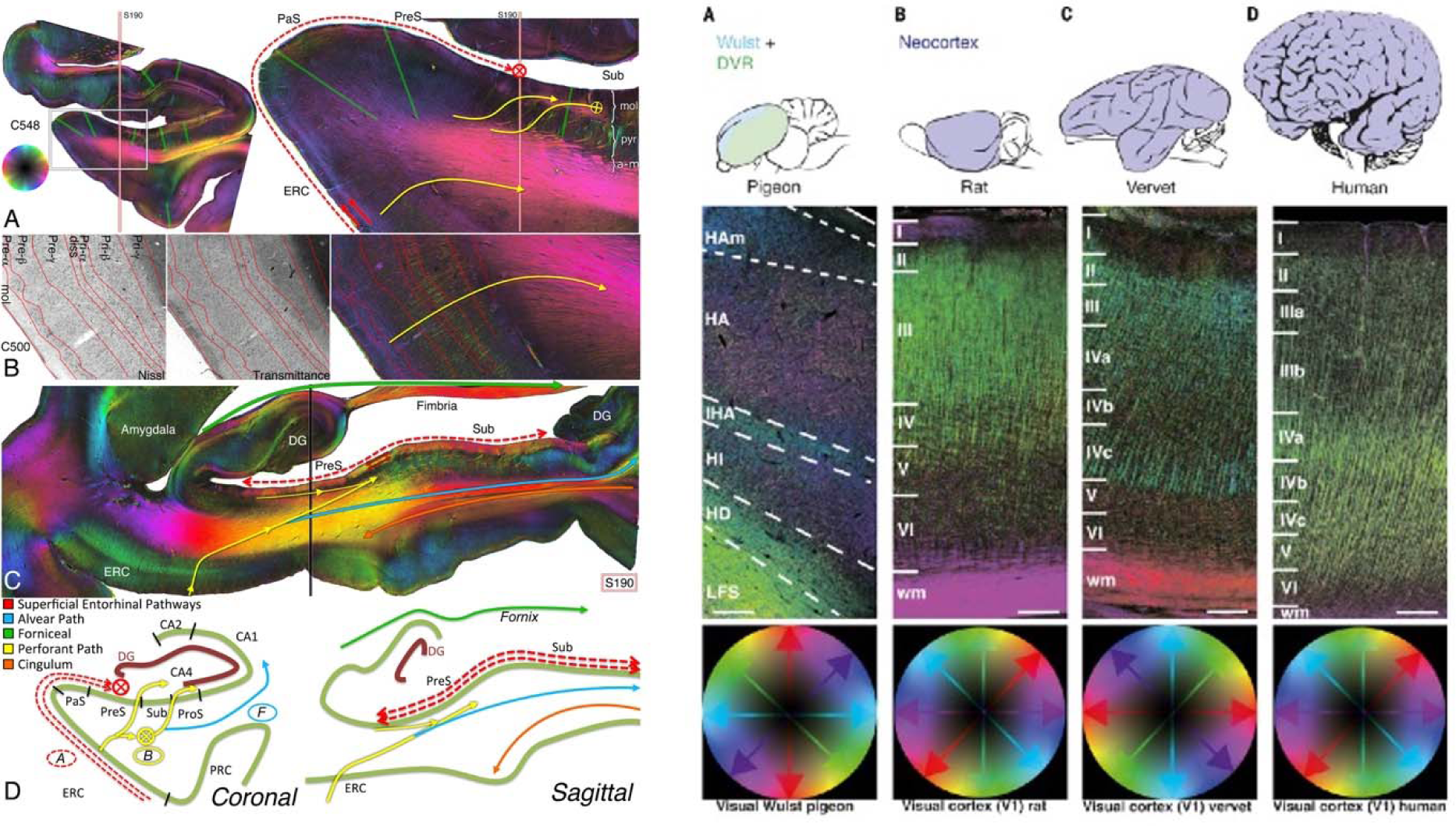
Fiber orientation maps acquired with 3D-PLI. **Left:** Entorhinal pathways and the angular bundle in the human hippocampus (reused from Zeineh et al., 2017). **Right:** 3D fiber architecture of the avian and mammalian primary visual regions (reused from Stacho et al., 2020). Fiber orientations are encoded in HSV color space.

*Optical coherence tomography (OCT)* is another approach to label-free imaging. Unlike the polarization microscopy methods described above, OCT does not operate in transmission mode, *i.e.,* the light does not go through a section of tissue. Instead, OCT relies on the back-scattering of light from a block of tissue, analogous to ultrasound technologies. As a result, it does not require the tissue to be sectioned before it is imaged. It uses optical interferometry to image depth-resolved tissue structures at resolutions in the order of 1–20 μm (Huang et al., 1991). After the superficial layer of the tissue block is imaged, it is sectioned to reveal and image the next layer. This technique has been successfully employed in various human brain applications both ex vivo and in vivo (Boppart et al., 1998; Böhringer et al., 2009; Assayag et al., 2013; Magnain et al., 2014, 2015, 2016, 2019).

*Polarization-sensitive OCT (PSOCT)* is a variation of OCT that was introduced by De Boer et al. (1997) to measure fiber orientations. It uses polarized light to probe birefringence. Wang et al. (2018) developed a fully *automatic, serial-sectioning PSOCT (as-PSOCT)* system for volumetric reconstruction of human brain samples at 3.5 *μm* in-plane and 50 *μm* through-plane voxel size. The as-PSOCT system is composed of a spectral domain PSOCT, motorized xyz translational stages enabling tile-wise imaging, and a vibratome to repeatedly remove a superficial layer slice of the formalin fixated tissue block upon completion of the full area scan. The implemented pipeline allows imaging and reconstruction of cm^3^ -sized tissue blocks over multiple days without the need for human intervention. The contrasts obtained with PSOCT include the light reflectivity provided by classical OCT, as well as additional contrasts derived from the polarization of light. The latter are the retardance, which is associated with myelin content, and the *en face* optic axis orientation, which is the orientation of fibers within the imaging plane. The use of PSOCT to image intricate fiber architectures in the brain has been demonstrated in mouse (Nakaji et al., 2008; Wang et al., 2011, 2016), rat (Wang et al., 2014a), and human (Wang et al., 2014b, 2018; Jones et al., 2020). Fig. 5 shows an example of PSOCT contrasts obtained from a human cerebellum.

**Figure 5.**
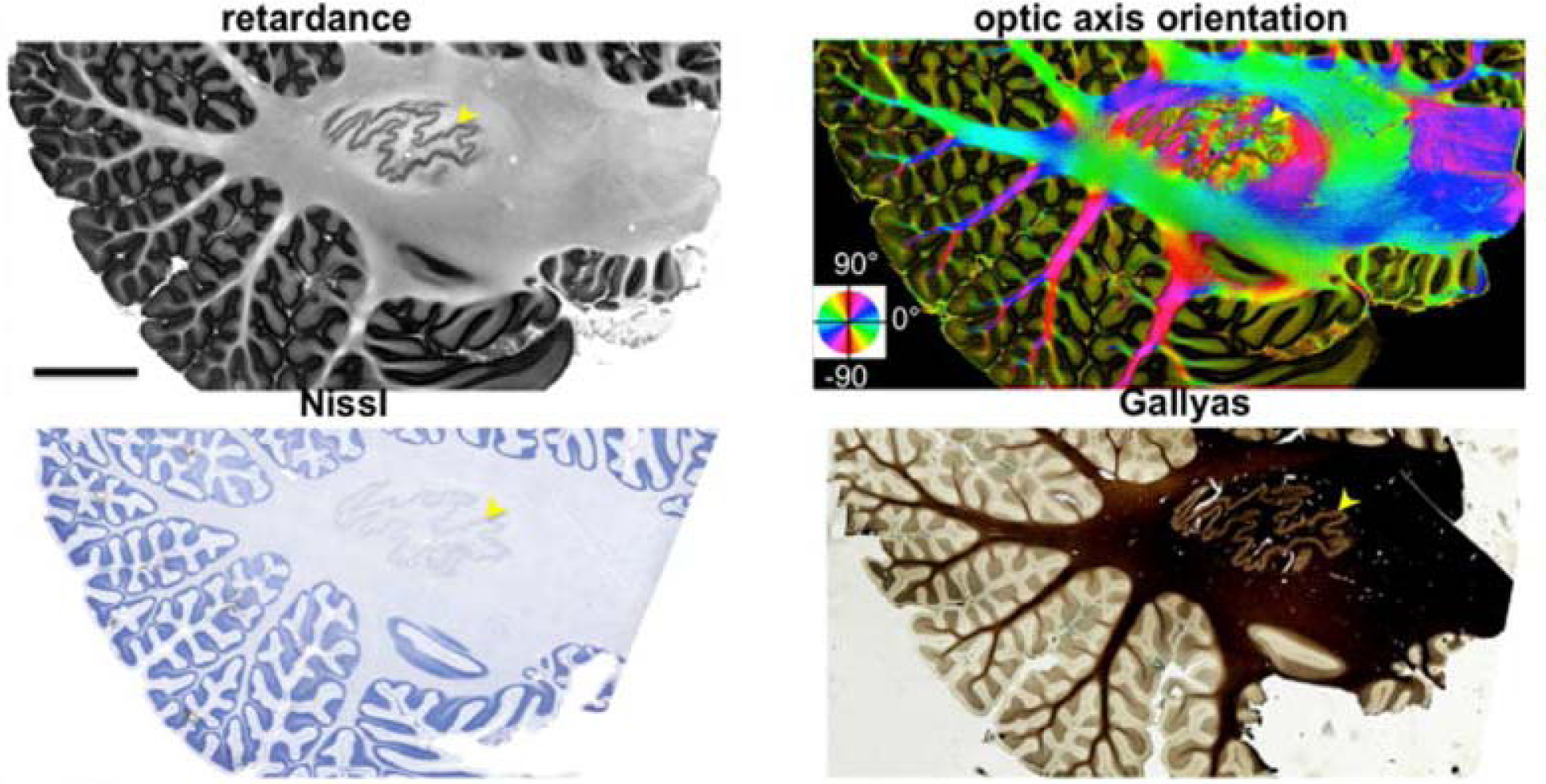
**Top:** PSOCT retardance and optic axis orientation maps of a 15 cm^2^ parasagittal section of the human cerebellum. **Bottom:** Nissl stain and Gallyas stain from the same sample. (Reused from Wang et al., 2018.)

Finally, an alternative to label-free methods is *tissue clearing,* followed by labeling, *e.g.,* with lipophilic dyes or immunohistochemistry of myelin-specific proteins, and imaging, *e.g.,* with *confocal fluorescence microscopy* (CFM) or *two-photon fluorescence microscopy* (TPFM). Tissue clearing can be performed with *Clear Lipid-exchanged Acrylamide-hybridized Rigid Imaging/Immunostaining/In situ hybridization-compatible Tissue-hYdrogel* (CLARITY; Chung et al., 2013). Clearing involves infusing the tissue with hydrogel, which is then polymerized to act as a support structure for the tissue. Lipids can then be removed from the tissue, rendering it optically transparent while preserving the rest of its biomolecular content. Cleared tissue samples can be imaged without sectioning and, consequently, without the need for tile stitching or slice alignment procedures. Tissue clearing methods are an active area of research, with ongoing efforts to make them applicable to larger samples and to the higher myelin density of the human brain (Tomer et al 2014; Yang et al., 2014; Costantini et al., 2015, 2019; Hou et al., 2015; Murray et al., 2015; Park et al., 2019; Zhao et al., 2020). After clearing, it is possible to perform consecutive rounds of staining with different fluorescent dyes on the same tissue sample with *System-Wide control of Interaction Time and kinetics of CHemicals* (SWITCH; Murray et al., 2015). For the purpose of comparison to dMRI orientation estimates, clarified tissue is typically stained for neurofilaments (Stolp et al., 2018; Leuze, et al 2021). As opposed to label-free techniques like PLI and PSOCT, which acquire direct measurements of axonal orientations, here the orientations have to be computed from the fluorescence microscopy images by structure tensor analysis, similarly to the conventional histological stains of section 5.1.2.

#### 5.2.2 Comparison to dMRI

Qualitative comparisons of fiber orientation maps obtained with PLI and dMRI from the same samples have been performed in the human brainstem (Henssen et al., 2019). A quantitative validation study compared the in-plane dispersion of fiber orientations in the human corpus callosum, as estimated from histology, PLI, and dMRI in the same samples (Mollink et al., 2017). The authors reported correlation coefficients of r = 0.79 between dMRI and histology, and r = 0.6 between dMRI and PLI.

A study that performed voxel-wise comparisons of optic axis orientation measurements from PSOCT and principal eigenvectors of tensors derived from dMRI showed low average angular error but high variability (5.4° ± 32.5°) in a human medulla oblongata (Wang et al., 2014b). A more recent study used PSOCT to evaluate the accuracy of dMRI orientation estimates obtained with different q-space sampling schemes (Cartesian, single- and multi-shell), spatial resolutions, and orientation reconstruction methods in human white-matter samples (Jones et al., 2020). The spatial resolution emerged as a key factor, with accuracy deteriorating as dMRI voxel size increased from 1 to 2 mm. In comparison, the benefit of increasing the number of directions from 128 to 514 and the maximum b-value from 12,000 to 40,000 was modest. The best- and worst-case mean angular errors, among all sampling schemes and orientation reconstruction methods at the native resolution of the dMRI data, were 8.3°/15.1° in single-fiber areas, 18.7°/35.7° in a fiber branching area, 20.2°/42.2° in an interdigitated crossing area, and 15.7°/32.7° in a separable crossing area. Thus the branching and interdigitated crossing were particularly challenging fiber configurations for dMRI.

Two studies have used CFM to image brain tissue processed with CLARITY, and compare FODs extracted with 3D structure tensor analysis (see section 5.1.2) to dMRI orientations in the same sample. The first study performed a qualitative comparison and showed moderate agreement between FODs from dMRI and those obtained from a variety of fluorescent dyes, including for neurofilaments (Stolp et al., 2018). The second study performed a quantitative comparison of FODs extracted from a neurofilament stain and dMRI orientations in the same human sample (Leuze et al., 2021). Angular errors of 19° ± 15° were reported between the principal fiber orientations obtained from CLARITY microscopy and dMRI, in a mainly single-fiber area. This error is about twice as big as the errors previously reported in single-fiber areas (see above and 5.1.2) using 2D structure tensor analysis on myelin stains (Leergaard et al., 2010; Choe et al., 2012; Seehaus et al., 2015), 3D structure tensor analysis on myelin stains (Schilling et al., 2016; Schilling et al., 2018) or 2D orientations measured with PSOCT (Jones et al., 2020).

#### 5.2.3 Limitations

Label-free optical imaging methods, like PLI and PSOCT, obtain direct measurements of axonal orientation angles. That is, no image processing operations like structure tensor analysis are necessary to estimate the fiber orientation vectors from the images. This is unlike methods that involve staining the tissue, whether conventionally (section 5.1.1) or after clearing with CLARITY. Removing both the possible tissue shrinkage sustained during conventional histological staining (see section 5.1.3) and the additional image processing step is the advantage of label-free methods like PLI and PSOCT.

However, like histological techniques, PLI requires tissue to be sectioned and mounted before imaging, which may introduce deformations. The importance of accurate cross-modal registration for mitigating histological distortions was discussed in section 5.1.3. The typical section thickness used for PLI is greater than that commonly used for conventional histology, and this may lead to somewhat less severe distortions. Custom registration frameworks have been developed for registering PLI to MRI (Ali et al., 2018).

The advantage of PSOCT in this regard is that it images the superficial layer of tissue *before* sectioning rather than after. Thus, the PSOCT orientation and retardance maps do not suffer from such deformations and do not require any registration between slices. However, PSOCT scanning is at present much slower, which limits the sample sizes that can be imaged in a practical amount of time (*e.g.,* 85 hours to scan a roughly 2 cm^3^ section with 3.9 μm in-plane resolution in Jones et al., 2020).

A possible concern that was discussed in the context of histological techniques (see section 5.1.3), and that also applies to label-free imaging methods when in-plane orientations are measured, is whether evaluating the accuracy of dMRI orientations in 2D results in any bias.

Encouragingly, the angular errors computed in single-fiber areas by PSOCT agree with those reported using conventional histological techniques either in 2D or in 3D (section 5.1.2). Furthermore, the in-plane angular errors between dMRI and PSOCT orientations were not found to be associated with the magnitude of the through-plane component of the dMRI orientations (Jones et al., 2020). One approach that has been proposed to infer through-plane fiber orientations with PSOCT is to apply structure tensor analysis along the through-plane dimension of the stacked retardance maps (Wang et al., 2018). Another possibility is to infer the through-plane orientation from measurements of in-plane orientations with two or more light incidence angles (Ugryumova et al., 2006; Ugryumova et al., 2009). In PLI, a forward model of birefringence and fiber orientation measurements from multiple oblique views is used to infer 3D orientations directly (Axer et al., 2011b; Schmitz et al., 2018). These techniques, however, have not yet been used in dMRI validation studies.

In principle, processing with CLARITY should allow tissue to be imaged intact, i.e., without sectioning. However, as the clearing process removes lipids from the myelin and membranes, the tissue becomes softer, deforms, and expands. Importantly, all MRI scanning must be completed before clearing, as the latter eliminates MR contrasts (Leuze et al., 2017). As a result, tissue deformation and expansion induced by clearing would make accurate alignment to dMRI challenging. As a result, studies that performed voxel-wise comparisons to dMRI used a passive form of CLARITY (Tomer et al 2014; Yang et al., 2014). This approach reduces tissue expansion but is slow and can thus only be applied to tissue sections (e.g., a 1 cm^2^ area with 500 μm thickness in Leuze et al., 2021). In Leuze et al (2017), a 315% increase in volume after one week of passive clearing was reported. Prior to microscopy, the clarified sections are placed in a refractive index matching solution that shrinks them back to approximately their initial size. After microscopy, affine registration is used to align sections to each other and/or to the dMRI volume (Stolp et al., 2018; Leuze et al., 2021).

A major challenge for all optical imaging techniques is scaling them to image the circuitry of the entire human brain at microscopic resolution. Cryo-sectioning and mounting sections with an area up to 15×10 cm^2^ and a thickness of 50 mm is well-known to result in section-unique, non-linear distortions. For PLI, where the brain is imaged after slicing, aligning large sections at the level of individual axons is difficult, even with extensive landmarks. For PSOCT, where each slice is cut after it is imaged, scaling up is a hardware problem, which will require the integration of a microtome that can handle an entire brain (frozen or embedded in solid materials) onto the OCT rig. For CLARITY processing, tissue volume is also not arbitrarily scalable in all directions, as the clearing solutions and dyes cannot penetrate an entire human brain. Deformations induced by clearing and sectioning once again make alignment of large slices at the level of individual axons a potentially insurmountable problem. Even if these technological hurdles are overcome in the coming years, optical imaging comes with staggering storage and computational requirements. For example, sampling a human brain volume of 1,200 cm^3^ with voxels sizes of 0.244 × 0.244 × 1 mm^3^ (TPFM), 1.3 × 1.3 × 60 mm^3^ (3D-PLI), or 3.5 × 3.5 × 15 mm^3^ (OCT) voxel sizes results in a total of 10^16^, 10^13^, or 10^12^ voxels, respectively. With multiple contrasts acquired and multiple subsequent processing steps, this leads to massive datasets that need to be transferred, stored, and analyzed. Access to supercomputing resources and specialized software will therefore be key for enabling fully digitized, high-throughput neuroanatomy at the microscale.

## 6. Validation of microstructure beyond fiber orientations

### 6.1 Microstructural modeling

Microstructural models aim to interrogate tissue properties such as axon diameter, density, packing and the degree of myelination, all of which are known to affect the diffusion of water molecules in the tissue (Beaulieu et al., 2002). To investigate how the microstructure varies between or along white-matter tracts, these models are fit to the dMRI signal on a voxel-by-voxel basis, and the parameters of interest are then typically either averaged over all voxels that intersect with a certain tract or averaged over consecutive cross-sections of the tract to generate an along-tract profile (Jones et al., 2005; Yushkevich et al., 2008; Colby et al., 2012; O’Donnell et al., 2009).

For details on the plethora of competing dMRI microstructural models and their relative strengths and weaknesses, we refer the reader to the many excellent reviews on the topic (*e.g.,* Jelescu and Budde, 2017; Novikov et al., 2018; Alexander et al., 2019; Dhital et al., 2019; Jelescu et al., 2020). Though the parameters of these models are sometimes given names such as “neurite density” or “axon diameter distribution”, they should often be thought of as sensitive “indices” rather than direct estimates of these tissue properties. This is because dMRI microstructural models make simplifying assumptions about complex tissue structure (Jelescu and Budde, 2017). These assumptions are often violated in real tissue but they are necessary to make the models parsimonious enough to be fit to in vivo dMRI data. As a result, microstructural measures are also in need of validation. In this section, we discuss some key post mortem imaging modalities that can be used for this purpose.

### 6.2 Inferring microstructure from post mortem tissue

For a rigorous comparison of microstructure between post mortem microscopy and MRI, both modalities must be acquired in the same tissue sample. There is considerable variety in the species that have been studied in the microstructure validation literature. Non-human primates demonstrate cortical folding and structural complexity closest to that of the human, but with a largely reduced brain volume, which makes dense microscopy imaging more practical. Rodent studies can benefit from a toolbox of interventions, including genetically engineered knockouts (Song et al., 2002; Nair et al., 2005; Kaden et al., 2016; Kelm et al., 2016) and induced diseases (Budde et al., 2008) or toxicity (e.g., in the cuprizone model of demyelination, see Torkildsen et al., 2013). These interventions can alter the tissue microstructure in controlled, well-specified ways. Furthermore, experimental animals typically undergo perfusion fixation at death, minimizing autolysis-related changes to the tissue microstructure such as the loosening and degradation of myelin (Hukkanen and Röyttä, 1987; Dyrby et al., 2018). In comparison, human tissue is immersion fixed. As a result, the postmortem interval, i.e., the time between death and fixation, can be relatively long, particularly for deep brain structures, leading to tissue degradation.

An important consideration for ex vivo validation of tissue microstructure is that microscopy images of postmortem, fixed, processed tissue, may not be a faithful representation of the in vivo structure (Howard et al., 2019). In addition, dMRI contrast differs ex vivo vs. in vivo. Formaldehyde fixation leads to the cross-linking of proteins, resulting in increased tissue rigidity and a reduction in diffusivity. This must be accounted for in diffusion microstructure models such as neurite orientation dispersion and density imaging (NODDI; Zhang et al., 2012), which assume fixed, pre-defined diffusivities for the tissue (Grussu et al., 2017). Furthermore, when applied to ex vivo data, microstructural models frequently include a non-diffusing “dot-compartment” (Grussu et al., 2017; Palombo et al., 2020; Veraart et al., 2020), which represents immobilized water in fixed tissue. Though most studies provide evidence for this compartment (Stanisz et al., 1997; Alexander et al., 2010; Panagiotaki et al., 2010; Kaden et al., 2016; Veraart et al., 2020), others do not (Palombo et al., 2020). Consequently, best practice can include fitting both the in vivo “default” and ex vivo “dot-variant” models, and comparing the relative quality of the model fits (Palombo et al., 2020), e.g., using the corrected Akaike information criterion (Burnham and Anderson, 2002).

### 6.3 Histological stains

Histological techniques were discussed in section 5.1 in the context of validating fiber orientations. Beyond the myelin and silver stains described in that section, there is a wide range of dyes and chemical compounds for the selective staining of other tissue constituents, such as cell bodies (e.g., NISSL stain) or glia (e.g. GFAP, which stains all glia, or IBA-1, which stains only activated microglia – a marker of inflammation). As each stain only informs on a subset of the tissue microstructure, sections can be counterstained (e.g., in H&E staining), or consecutive slices can be processed for different stains, which together build a more complete characterization of tissue composition.

Quantitative comparisons of histology and dMRI microstructural parameters may require cell counting, characterizing the shapes and sizes of cells, extracting staining densities, or calculating the fraction of stained pixels over a local neighborhood. When stains such as DAB do not follow the Beer-Lambert law, the relationship between the stain density and chromogen concentration in a pixel is not linear. Consequently, the stain density is often considered only a proxy or semi-quantitative histology metric (Tolcos et al., 2016).

Histology metrics have been compared to measures from most common dMRI microstructural models. Below we provide only a few examples.

#### Anisotropy and diffusivity

At the timescales of typical diffusion experiments, myelinated fibers appear approximately impermeable to water. Consequently, histological estimates of myelin density are frequently compared with parameters related to diffusion anisotropy. Meta-analyses of studies that have compared myelin content from histology to parameters from diffusion tensor imaging (DTI; Basser et al., 1994) reveal that myelin content correlates positively with fractional anisotropy (FA) and negatively with radial diffusivity, but not with axial diffusivity or mean diffusivity (Mancini et al., 2020; Lazari and Lipp, 2021). As an alternative to myelin content, a tissue anisotropy index computed by structure-tensor analysis of DiI-stained histological sections has also been found to correlate with FA in rat (Budde and Frank, 2012) and macaque (Khan et al., 2015).

#### Axon calibre

When imaged at high resolution (∼10 nm in-plane), neurofilament-stained sections can validate estimates of axon radii (Veraart et al., 2020). Using this method in fixed rat tissue, Veraart et al. (2020) estimated effective axon diameters of ∼0.8-1.2 μm, which were closely aligned to, though slightly lower than, those estimated from a dMRI model using ultra-strong gradients (∼1-1.4 μm).

#### Neurite density

Histological myelin is often used as a proxy for neurite density, and has been found to correlate positively with the intra-axonal signal fraction estimated from various multi-compartment diffusion models (Jespersen et al., 2010; Grussu et al., 2017).

#### Cell density

NISSL-stained histology sections map the density of cell bodies across the brain. In the cortex, NISSL densities qualitatively mirror recent diffusion maps of cell soma (Palombo et al., 2020).

#### Glia density

Though glia morphologies are exquisitely characterized by histology, the relationship between glia and diffusion parameters remains relatively unexplored. Recently, increased density of activated microglia, a marker of inflammation, has been related to reduced FA in traumatic brain injury (Soni et al., 2020) and, intriguingly, the NODDI dispersion parameter (Yi et al., 2019), which Yi et al. argue is a marker for greater hindered diffusion in the extra-cellular space.

### 6.4 Optical imaging

Label-free optical imaging methods, and particularly PLI and PSOCT, were discussed in section 5.2, in the context of measuring fiber orientations. These techniques can also quantify the amount of myelin per pixel (Axer et al., 2011b; Wang et al., 2018; Mollink et al., 2019; see also, Fig. X2). A study that compared retardance as measured with PSOCT and FA computed from DTI data in the same human medulla oblongata sample reported correlation as high as r = 0.9 (Wang et al., 2014b). Another study showed moderate correlations between reflectivity and attenuation as measured with (non-PS) OCT and FA in mouse brain (Lefebvre et al., 2017).

Fluorescence microscopy, where fluorescent proteins or dyes label specific structures of interest, has also been used to validate dMRI microstructural parameters. This technique can provide quantitative estimates of a given structure within the tissue, as the fluorescence signals are often linearly related to the amount of fluorophore on the tissue. For example, a study using CFM in the spinal cord of a pre- and post-symptomatic ALS mouse model found that decreases in FA and axial diffusivity and increases in radial diffusivity correlated with changes in axonal fluorescence intensity and membrane cellular markers (Gatto et al., 2018).

The use of fluorescence dyes on clarified tissue was also discussed in section 5.2. Clarified mouse brains, imaged with CFM or TPFM, have been used to validate DTI measures in. One study showed positive correlations of neurite density and FA that were strong (r = 0.87) in medial lemniscus and less strong (r = 0.51) in caudate nucleus and putamen (Kamagata et al., 2016). Another study reported that the density of myelin basic protein immunofluorescence correlated strongly with FA and, in some cases, with radial diffusivity (Chang et al., 2017). A study in mouse hippocampus found staining for cell nuclei to be correlated with mean, radial, and axial diffusivity; neurofilament staining to be correlated with mean and radial diffusivity; and astrocyte staining to be correlated with FA (Stolp et al., 2018).

### 6.5 Electron microscopy

Electron microscopy (EM) can resolve nanometer-scale structures to characterize the detailed morphology of neurons and glia, or fine anatomical structures such as layers of the myelin lamella, synapses on post-synaptic neurons, microtubules in the cytoskeleton or mitochondria within the axoplasm (Kasthuri et al., 2015). For EM imaging, small tissue blocks are stained with heavy metals such as osmium, which shows excellent contrast for cellular membranes (Salo et al., 2021).

Methods for EM can vary both in terms of tissue processing (e.g., tape-based or block-face) and in terms of imaging techniques (e.g., back-scattered or transmission EM) (Lichtman et al., 2014). In serial, block-face scanning EM (SBEM), the top face of the sample is first imaged (using back-scattered electron detection) and then sectioned in situ, using a microtome or focused ion beam. The process is then repeated to produce a series of well-aligned images. Alternatively, using the tape-based method (Kasthuri et al., 2015), the tissue block is first sectioned using an ultramicrotome and mounted onto tape, after which the sections are imaged serially by either transmission or back-scattered EM. Where the SBEM images exhibit little deformation or misalignment, tape-based serial sections can be rotated, stretched, or otherwise misaligned with respect to each other, making reconstruction of the 3D volume more challenging. This is counterbalanced by the more limited lateral resolution of SBEM compared to transmission EM.

A key challenge in the analysis of EM data is the identification and segmentation of the various tissue types from the complex texture of the reconstructed 3D volume. The segmentation task is challenging for several reasons (Lichtman et al., 2014): *(i)* the stain is non-specific, hence different cell types must then be identified purely based on their morphology and/or localization with respect to other structures, *(ii)* the in-plane and through-plane resolution of the non-isotropic images can differ by an order of magnitude (e.g. 3×3×30nm), *(iii)* the cells have complex morphology and are often intertwined, and *(iv)* cellular processes often branch or fork, in which case two apparently separate structures in one image can in fact be part of the same cell. Automated cell segmentation has greatly benefited from recent advances in machine learning approaches (Abdollahzadeh et al., 2021), but some manual correction is often required for high precision (Kasthuri et al., 2015).

The acquisition time and data storage requirements of EM constrain the field of view that can be imaged. If a single cubic millimeter of tissue, *i.e.,* one MRI voxel, were imaged with EM at nanometer resolution, it would yield over 2TB of data, and would likely take several years on even some of the most advanced imaging systems (Lichtman et al., 2014). Consequently, nanometer EM techniques are limited to characterizing small tissue blocks (50-100 μm^3^), which are sparsely sampled throughout the brain. If the small EM block is not representative of the larger MRI voxel from which it is sampled, this may bias the comparison.

Tissue shrinkage is another concern for the use of EM in microstructure validation. Unless a specific extra-cellular space preserving method is adopted (Korogod et al., 2015; Pallotto et al., 2015), EM images typically suggest little or no extra-cellular space, precluding EM-based validation of this compartment. In comparison, axonal volumes appear relatively unaffected (Korogod et al., 2015; Jelescu and Budde, 2017).

Nonetheless, EM has been used fairly extensively to validate, challenge, and inspire biophysical diffusion modelling of tissue.

#### Axon undulation

The nanoscale resolution of 3D EM allows the complex, inhomogeneous morphology of axons to be visualized. The axons undulate through space and their diameter and degree of myelination varies along the axon length (Lee et al., 2019). Though many common diffusion models assume that axons are straight cylinders of fixed diameter, the effect of complex axon morphologies on the diffusion signal is an area of active investigation (Budde and Frank, 2010; Nilsson et al., 2012; Fieremans et al., 2016; Ginsburger et al., 2018; Lee et al., 2019, 2020a,b; Brabec et al., 2020; Jelescu et al., 2020).

#### Axon calibre

By fitting a circle or ellipse to the circumference of axons, EM data can be used to estimate axon diameters, or the axon diameter distribution over a given area. The axon diameters estimated from EM data range from 0.1-10 μm, with the vast majority of axons having diameters below 3 μm (Aboitiz et al., 1992; Lee et al., 2019). However, dMRI estimates of axon diameter, e.g., from the composite hindered and restricted model of diffusion (CHARMED; Assaf and Basser, 2005), AxCaliber (Assaf et al., 2008), or ActiveAx (Alexander et al., 2010), can be up to an order of magnitude larger (Alexander et al., 2010). This discrepancy is now thought to be related to the dMRI signal from clinical-grade scanners lacking sensitivity to the intra-axonal space (Jelescu et al., 2020). Nonetheless, axon diameters obtained from diffusion models are able to replicate known variations across the brain (Fan et al., 2020), such as the low-high-low pattern of axon diameter across the corpus callosum, as was previously characterized in EM data by Aboitiz et al. (1992).

#### G-ratio

EM data can also be used to estimate and validate estimates of the g-ratio, i.e., the ratio of the inner and outer (axon + myelin) diameter of the axon (Stikov et al., 2015).

#### Intra-axonal volume fraction

EM data is also used to validate estimates of the intra-axonal signal fraction from multi-compartment diffusion models (Jespersen et al., 2010; Kelm et al., 2016). Here, EM estimates of the relative fractions of the intra-axonal, myelin, and extra-axonal space can be converted to MR signal fractions by accounting for the *T*_2_ properties of the various compartments. This includes correcting for the substantial fraction of myelin, which is “MR-invisible” at the echo times of typical diffusion experiments due to its ultra-short *T*_2_ (∼10ms, see Mackay et al., 1994). When corrected, EM estimates of 30-50% intra-axonal space in the white matter (Xu et al, 2014; Stikov et al., 2015; Jelescu et al., 2016; Lampinen et al., 2019) translate to MR-visible fractions of 40-60%, as is found in many diffusion models (Jelescu and Budde, 2017).

### 6.6 X-ray microcomputed tomography

Micron-scale synchrotron x-ray computed tomography (microCT) is a non-destructive, 3D imaging technique that can provide micron resolution images of thick tissue blocks (∼1cm cubed) (Foxley et al., 2020). Here, osmium-stained tissue is rotated through a synchrotron x-ray beam, after which a high-resolution 3D volume is reconstructed from a series of 2D x-ray back-projections. MicroCT of neural tissue can capture complex, 3D cellular morphologies (Andersson et al., 2021), as well as detailed, long range axonal projections (Foxley et al., 2020), making it a valuable, up-and-coming tool for microstructure validation.

## 7. Discussion

In the previous sections we described a variety of methods for validating dMRI tractography, fiber orientations, and other microstructural properties of axon bundles in post mortem tissue. As should be clear by now, each of these methods has its own potential sources of errors and uncertainty. When faced with this fact, one may ask, “Is this really ground truth?” There are several important considerations here. First, these methods are not meant as push-button solutions for obtaining a correct answer. They should always be coupled with neuroanatomical expertise. The experienced anatomist can interpret the results of a validation study, place them in the context of centuries of knowledge on neuroanatomy, and ultimately evaluate the quality of the validation dataset itself. Second, small errors at the microscopic scale where these methods operate can still yield accurate results at the mesoscopic scale of dMRI. Third, there is great value in obtaining converging information from multiple sources of validation data. The fact that none of these validation methods are error-free or can provide ground truth on every cell in the brain does not mean that we should use none of these methods but that we should use many of them.

Efforts on post mortem validation may appear to progress slowly; data sets are small in comparison to *in vivo* neuroimaging, and require time-consuming data collection and post-processing. Nonetheless, we present here a growing body of knowledge, with several implications emerging for how dMRI data should be acquired and analyzed. For example, there is now accumulating evidence from tracer validation studies that probabilistic tractography has greater anatomical accuracy than deterministic tractography. This comes from studies that compared area-to-area connectivity matrices between dMRI tractography and existing tracer databases (Delettre et al., 2019; Girard et al., 2020), as well as studies that performed voxel-wise comparisons of dMRI tractography and tracer injections in the same NHP brains (Grisot et al., 2021; Maffei et al., 2020, 2021).

Furthermore, evidence on the limited benefit of very high *b*-values on the accuracy of ODFs and tractography is emerging from several independent sources, including histological staining (Schilling et al., 2018), PSOCT (Jones et al., 2020), and anatomic tracing (Ambrosen et al., 2020; Grisot et al., 2021). This does not necessarily mean that very high *b*-values do not contain useful information on fiber architectures. It may mean that existing methods for ODF reconstruction and tractography are not designed to take advantage of this information. This topic merits further investigation in the future. The same is true of converging evidence from PSOCT (Jones et al., 2020) and anatomic tracing (Grisot et al., 2021; Maffei et al., 2021) on the importance of resolving fiber configurations other than crossing, such as branching or turning, and the role that spatial resolution might play towards this end.

The studies that we have reviewed here offer a glimpse of the knowledge that can be gained by bridging the gap between whole-brain, mesoscopic-resolution MRI and microscopic-resolution imaging of smaller samples, ideally from the same brain. New technologies developed by the Connectome 2.0 project, discussed elsewhere in this issue (Huang et al., TBD), aim to push spatial resolution for both *in vivo* and *ex vivo* dMRI to the sub-mm level, thus narrowing this gap. Advanced computational methods will be necessary to improve alignment across modalities and overcome the differences in shape, dimensions, physical and morphological characteristics of the tissue at every step of data acquisition. A nested connectome reconstruction across scales, that combines the complementary virtues of many imaging techniques and utilizes machine learning approaches to predict missing information, will be advantageous for this purpose. For example, recent work used deep neural networks to predict fluorescence images of diverse cell and tissue structures from measurements of density, anisotropy, and orientation acquired with label-free imaging (Guo et al., 2020). Other examples include joint modeling of microscopy and dMRI, *e.g.,* to overcome the degeneracy between fiber dispersion and radial diffusion in dMRI models (Howard et al., 2019), and training neural networks on paired optical imaging and dMRI data to infer complex fiber architectures directly from dMRI (Yendiki et al., 2020). Ultimately, the next frontier for the microscopic imaging modalities described in this review will be to use them not only to validate existing computational tools for inferring brain circuitry from mesoscopic dMRI, but to engineer the next generation of such tools.

## Acknowledgements

A. Yendiki was supported by NIH grants R01-EB021265, R01-NS119911, P50-MH106435, and U01-EB026996. M. Aggarwal was supported by NIH grant R01-AG057991. M. Axer was supported by the European Union’s Horizon 2020 Framework Programme for Research and Innovation under the Specific Grant Agreement No. 945539 (“Human Brain Project” SGA3). A.F.D. Howard and the Wellcome Centre for Integrative Neuroimaging were supported by the Wellcome Trust (grants WT202788/Z/16/A and 203139/Z/16/Z). S.N. Haber was supported by NIH grants R01-MH045573 and P50-MH106435.

